# Host genomic influence on bacterial composition in the switchgrass rhizosphere

**DOI:** 10.1101/2021.09.01.458593

**Authors:** Jeremy Sutherland, Terrence Bell, Ryan V. Trexler, John E. Carlson, Jesse R. Lasky

**Affiliations:** Department of Plant Pathology and Environmental Microbiology, The Pennsylvania State University, University Park, PA, USA; Intercollege Graduate Degree Program in Bioinformatics and Genomics, The Pennsylvania State University, University Park, PA, USA; Intercollege Graduate Degree Program in Ecology, The Pennsylvania State University, University Park, PA, USA; Department of Ecosystem Science and Management, The Pennsylvania State University, University Park, PA, USA; Department of Biology, The Pennsylvania State University, University Park, PA, USA

**Author notes:** Corresponding author contact: 1-814-863-5318.

**Keywords:** switchgrass, rhizosphere, microbiome, host genomic influence, heritability

## Abstract

Host genetic variation can shape the diversity and composition of associated microbiomes, which may reciprocally influence host traits and performance. While the genetic basis of phenotypic diversity of plant populations in nature has been studied, comparatively little research has investigated the genetics of host effects on their associated microbiomes. Switchgrass (*Panicum virgatum*) is a highly outcrossing, perennial, grass species with substantial locally adaptive diversity across its native North American range. Here, we compared 383 switchgrass accessions in a common garden to determine the host genotypic influence on rhizosphere bacterial composition. We hypothesized that the composition and diversity of rhizosphere bacterial assemblages would differentiate due to genotypic differences between hosts (potentially due to root phenotypes and associated life history variation). We observed higher alpha diversity of bacteria associated with upland ecotypes and tetraploids, compared to lowland ecotypes and octoploids, respectively. Alpha diversity correlated negatively with flowering time and plant height, indicating that bacterial composition varies along switchgrass life history axes. Narrow-sense heritability (*h*^2^) of the relative abundance of twenty-one core bacterial families was observed. Overall compositional differences among tetraploids, due to genetic variation, supports wide-spread genotypic influence on the rhizosphere microbiome. Lastly, a genome-wide association study identified 1,861 single-nucleotide polymorphisms associated with 110 families and genes containing them related to potential regulatory functions. Our findings suggest that switchgrass genomic and life-history variation influences bacterial composition in the rhizosphere, potentially due to host adaptation to local environments.

## Introduction

Macroscopic organisms host diverse microbiomes, which often differ substantially among individuals (Smith et al., 2015; Trivedi et al., 2020; Wilson et al., 2020). Some of the proximate causes of this variation in microbiome composition arise from differences between host environments and traits, the latter of which can be divided into genetic and plastic components. Host genetic differences can affect a wide range of traits that influence associated microbiome composition, including traits that determine the supply of resources used by microbes or traits that control environmental variables affecting microbial fitness in other ways (e.g. pH) (Lebeis et al., 2015; Sasse et al., 2018). These host genetic influences on microbiomes may arise from different evolutionary processes. For example, host traits that affect microbiome composition may evolve neutrally or in response to non-microbial components of the local environment (Zeng et al., 2015). Alternatively, host trait variation could have evolved as a direct response to beneficial interactions with microbes in the environment (Gould et al., 2018). In the latter case, hosts could either influence the abundance of microbes that are simply present in the environment, and functionally important to the host, or hosts could form species or strain-specific symbiotic relationships where the host depends on and influences the abundance of those taxa for reproductive success (Parker et al., 2006).

Because of environmental and dispersal limitations, the same microbes may not be present across the entire range of a host, although functionally similar ones might be. It has been shown that reciprocal interactions between host immune systems and host-associated microorganisms can lead to preferential recruitment of beneficial microbial types and repel pathogenic types, to varying degrees (Hacquard et al., 2017; Jones et al., 2019). This suggests that hosts might (directly or indirectly) select for functional microbial traits from certain lineages. Several studies have observed the presence of ‘core microbiota’ – persistent members of the microbiome that appear across a large portion of a host’s population or species (Risely, 2020), but it is often unclear whether these microorganisms are functionally important or commensal to the host. Yet, because microbial traits are often phylogenetically conserved and host selection of microbes can occur in related microbial lineages (Lemanceau et al., 2017; Martiny et al., 2015), a core microbiome might be preserved for higher taxonomic levels but undetectable for lower ones, at the same occupancy frequency threshold. The study of diverse host genotypes and phenotypes in common environments could reveal the genetic and host physiological drivers of variation in associated microbiomes and, potentially, types of microbes that are functionally important to the host.

The demonstrated influence of host trait variation on microbial assemblages (Hestrin et al., 2021; Jones et al., 2019; Wagner et al., 2016) suggests that microbiome composition can be treated as an extension of host phenotype, one that may be under selective pressure by both the host and the environment (Bordenstein & Theis, 2015; Hunter, 2018; Moran & Sloan, 2015; Wagner et al., 2016; Whitham et al., 2003). Genome-wide association studies (GWAS) have shown that the relationship between hosts and specific microbial taxa can be linked to single-nucleotide polymorphisms (SNPs) (Beilsmith et al., 2019). However, at the microbiome level, it becomes challenging to directly link host genotype and microbial composition, due to the high diversity and dimensionality of microbiomes (Wray et al., 2013). Common garden experiments and genetic mapping offer a window into host genetic impacts on host-associated microbial composition. The resolution of association mapping can be improved by exploiting high levels of genomic diversity within and between populations (Holland, 2007).

Switchgrass (*Panicum virgatum)* is a C4 perennial grass species that has been extensively studied as a bioenergy crop since 2005 (Bouton, 2007; McLaughlin & Kszos, 2005; Parrish & Fike, 2005; Sanderson et al., 2006). Unlike first generation biofuels, switchgrass can be grown on marginal land with minimal inputs while also providing ecological services (e.g. wildlife habitat and soil erosion mitigation) (Werling et al., 2014). In addition to its ecological and agricultural value, switchgrass provides an ideal system for studying the influence of host genotypic diversity on rhizosphere bacterial composition (i.e., bacteria living on or in close proximity to the root system of the plant) because of the high degree of genotypic diversity in the species.

Switchgrass is primarily an outcrossing species with high genetic diversity within populations, showing evidence of inbreeding depression, which increases heterozygosity in the genome (Sharma et al., 2012). The species is comprised of two locally-adapted ecotypes, upland and lowland, based on differences in phenotype, physiology, and habitat (Das et al., 2004). Separate migration events fostered by interglacial periods, in addition to gene duplication through polyploidization, have also played a major role in shaping the genomic diversity of switchgrass (Casler et al., 2015). Lowland switchgrass is primarily allotetraploid (4X: n = 36) and exhibits disomic inheritance – chromosome inheritance that follows diploid association patterns. Upland accessions tend to vary in their ploidy level (4X: n = 36, 6X: n = 54, 8X: n = 72) and exhibit widespread aneuploidy between and within localized populations (Costich et al., 2010). Additionally, populations of switchgrass are adapted to local climate conditions and linkage mapping suggests a polygenic basis to this local adaptation (Lowry et al., 2019). Here, we studied genotypes largely from the geographically Northeastern, upper Southern, and Midwestern United States that can be genetically clustered into five groups (roughly North, South, East, West, and Northeast, with some geographic overlap) (Supplemental Figure 1) (Lu et al., 2013). Genetic variation in switchgrass has been linked to measures of plant fitness and performance, including biomass production (Lowry et al., 2019) and soil carbon inputs (Adkins et al., 2016; Stewart et al., 2017). While substantial research has explored the switchgrass microbiome, key gaps remain, including the functional role of non-fungal microorganisms and host mechanisms that underly switchgrass– microbiome interactions (Hestrin et al., 2021).

Vascular plants influence microbes in the rhizosphere through rhizodeposits (Dennis et al., 2010) and carbon-based root exudates (Compant et al., 2010; Hu et al., 2018; Yu & Hochholdinger, 2018). Root exudates provide signaling and substrate compounds for beneficial microbial recruitment (Carvalhais et al., 2013; Lakshmanan et al., 2012), and antibiotic compounds that defensively modulate the microbiome composition against pathogenic taxa (Baetz & Martinoia, 2014; Lebeis et al., 2015). When available, root exudates potentially represent a larger portion of carbon in the soil compared to other carbon inputs from the environment (e.g. leaf litter) (Heijboer et al., 2018). Switchgrass can benefit from a range of microbes that are influenced by root exudates (Hestrin et al., 2021).

In this study, we analyzed bacterial 16S rRNA gene sequences from total DNA extracted from rhizosphere soil samples for 383 switchgrass accessions from 63 local populations planted within a common garden in Ithaca, NY, USA. Our aim was to better understand the influence of switchgrass genotype on the composition of bacteria in the rhizosphere. We hypothesized that overall bacterial composition and diversity within the growing site would differentiate due to host genetic variation along the three dimensions of switchgrass genetic diversity (ecotype, ploidy level, and genetic cluster), as is true for multiple fitness-related traits (Lovell et al., 2021). Separately, bacteria experience higher rates of gene loss, horizontal gene transfer, and shorter generation times compared to their hosts. So, microbial specialization on a given host can be evolutionarily fast compared to changes in the plant community (Bever et al., 2012). Given that the evolutionary response rate is mismatched between switchgrass genotypes and their microbiomes, we sought to determine the lowest taxonomic level for which a potential host influence could be detected. We show that switchgrass genotype influences bacteria in the rhizosphere at, at least, the family level. Our findings demonstrate how plant host genotype can influence host-associated microbiomes, the taxonomic level at which the microbiome is influenced, and the host genetic architecture involved.

### Materials and Methods Switchgrass Germplasm Data

Previously, exome capture sequencing was used to characterize variation in the genic regions of the genome from the Northern Switchgrass Association Panel used in this study (*n* = 537) (Evans et al., 2018). For this study, the HapMapv2 matrix was obtained from the Dryad Digital Repository (10.5061/dryad.mp6cp) in 2019. Several modifications were made to the original dataset. The dataset was modified in *python* (G. Van Rossum & Drake, 2009) to reflect heterozygote positions more conservatively than the original, unfiltered, dataset, such that heterozygote positions with read count ratios greater than 0.75 or less than 0.25 were called homozygous relative to the appropriate allele (e.g. A(20)/T(1) = A). SNPs with a minor-allele frequency (MAF) below 0.05 were excluded, resulting in 103,776 SNPs for all genotypes studied here (*n* = 362) spanning eighteen chromosomes: Chr01-09 (subgenomes a and b) of the v1.1 *P. virgatum* reference genome (www.phytozome.net) and fifteen unanchored scaffolds (Evans et al., 2018). The unanchored scaffolds of Evans et al. (2018) were renamed to the corresponding contig names within the v1.1 reference genome using *python*. Phylogenetic tree generated in Tassel v5 (Bradbury et al., 2007).

We compared bacterial composition to published data on several important switchgrass functional and life history traits, using data from 2009, 2010, and 2011 field seasons at the common garden in Ithaca, NY (genetic variation in these traits is largely consistent from year to year) (Lipka et al., 2014). The 2011 measurements for anthesis date and full plant height were used to identify correlations with rhizosphere alpha diversity present in 2016.

### Rhizosphere sampling of switchgrass accessions

The original switchgrass association panel consisted of 3-10 clonally propagated individuals each from 66 discrete switchgrass local populations grown from seed in the greenhouse of the USDA-ARS Dairy Forage Research Center in Madison WI in 2007 (Lu et al., 2013). The accessions were sourced from much of the natural range of switchgrass in the eastern and midwestern United States. Young plants were deposited in the common garden in Ithaca, NY, USA in 2008. In 2016, plugs were collected for transplanting from 525 cloned accessions, representing all but two (perished) original populations. Prior to replanting the plugs, soil samples were collected into a polythene bag from each of the switchgrass accessions by vigorously shaking off soil attached to the roots. The bags with soil and fine roots were initially stored at 4°C while being processed - each soil sample was thoroughly mixed into a uniform distribution and then transferred to 50 ml conical cryogenic tubes and stored at −80°C prior to soil DNA extraction.

### Soil DNA extractions

To determine the composition of bacteria established in the switchgrass rhizosphere, we selected soil samples from 383 of the 525 cloned switchgrass accessions, based on a phylogenetically determined set of clones from 63 of the local populations in the association panel. Total DNA was extracted from ~300 mg of soil using the NucleoSpin Soil 96 kit (Macherey-Nagel, Düren, Germany). Lysis was performed using the Buffer SL with Enhancer SX, and was performed on the FastPrep 24 homogenizer (MP Biomedicals, Santa Ana, CA, USA) at 4.0 m/s for 30 s.

### Amplicon Sequencing

Briefly, amplicons targeting the V3-V4 region of the 16S rRNA gene were generated from all 383 samples using universal bacterial primers 515F (5′-GTGYCAGCMGCCGCGGTAA-3′) and 806R (5′-GGACTACNVGGGTWTCTAAT-3′) (Apprill, Mcnally, Parsons, & Weber, 2015; Parada, Needham, & Fuhrman, 2016) with overhangs for attaching barcodes and standard Illumina overhang adaptors in a second PCR step (full protocol outlined in the supplemental methods of Trexler & Bell (2019)). Sequencing was performed on an Illumina MiSeq using the 2 × 250 cycle v2 kit. Raw reads are available through NCBI SRA under project number PRJNA689762.

### Amplicon sequence analysis pipeline

The raw 16S rRNA gene sequences were analyzed using an adapted version of the *dada2* pipeline (Callahan et al., 2016). *dada2* was used to process the raw sequences into Amplicon Sequence Variants (ASVs). Reads were truncated above 240 bp and below 160 bp. Sequences with any missing reads after truncation were discarded. Reads were then truncated at the first instance of a quality score less than 2. After truncation, reads that matched against the phiX Genome were discarded. Reads with higher than two “expected errors” were also discarded. Expected errors are calculated from the nominal definition of the quality score: EE = sum(10^(-Q/10)^). Filtered sequences were used to determine the error rate using the *dada2* function: learnErrors(). The filtered sequences were then processed with the core sample inference algorithm, dada(), incorporating the learned error rates. Forward and reverse sequence reads were then merged. A sequence table was made and chimeras were removed using the “consensus” method. Taxonomic assignments for the ASVs were defined against the SILVA 138 ribosomal RNA gene database (Quast et al., n.d.) using DECIPHER v2.14.0 (Wright, n.d.). The sequence variant table and taxonomy table were exported for downstream processing in *phyloseq* (McMurdie & Holmes, 2013).

In *phyloseq*, ASVs designated “NA” at the phylum level, non-Bacterial entries at the domain level, and those associated to chloroplasts and mitochondria were removed from the taxonomy file before further processing.

Finally, we also removed samples with no ASVs remaining after pruning. Rarefying samples has been shown to effectively reduce false discovery rates when there are large differences between the average sample library size (Weiss et al., 2017). Our data was characteristic of this scenario. Therefore, we rarefied samples to 1000 sequences, to retain as many samples as possible while also maintaining a sufficient sampling depth to detect the most prevalent taxa comprising the switchgrass core microbiome (See below).

### Alpha diversity analysis

Using the rarefied ASV dataset, we generated alpha diversity metrics using *phyloseq* (McMurdie & Holmes, 2013). Statistical significance was calculated using the vegan 2.5.6 package (Oksanen et al., 2019), and other core functions in R version 3.6.3 (Ihaka & Gentleman, 1996). The Chao1 and Shannon diversity indices were calculated using the estimate_richness() function in *phyloseq* and then used to test for differences in alpha diversity between genotypes differing in ecotype, ploidy level, and genetic cluster groups using an analysis of variance (ANOVA) (R version 3.6.3). Correlations between diversity metrics, anthesis date and full plant height were generated using the Pearson correlation test.

### Beta diversity and core microbiome analysis

Bacterial diversity data rarely conform to the assumptions of MANOVA-like procedures, largely due to inflated zero values among rare taxa that skew the distribution, thus non-parametric methods based on permutation tests are preferred (M. J. Anderson, 2001). A non-parametric, one-way analysis of variance (NPMANOVA) is also tolerant of non-independent observations (e.g. microbe-microbe interactions). We therefore calculated an NPMANOVA using the *adonis2* function in vegan to test for significant differences between the ecotype, ploidy level, and genetic cluster groups on the overall rhizosphere microbiome composition using pairwise Bray-Curtis distances at the ASV level.

The ‘core microbiome’ can be defined as microbial taxa that associate with a host above a particular occupancy frequency threshold (i.e. their prevalence). Biological justifications for such thresholds tend to be subjective, often ranging between 30% and 95% (Risely, 2020). However, core taxa prevalence within a population could be a reflection of an environmental response by the host due to local adaptation, resulting in higher intergroup prevalence of certain ‘core taxa’ relative to the whole population. Thus, we determined two core microbiomes: A high-fidelity core (prevalence >= 90%), to determine taxa that reliably associate with the vast majority of our panel, and a low-fidelity core (prevalence >= 10%), to study host genetic variation effects on bacterial composition. Taxa present within our rarefied dataset were initially agglomerated using the tax_glom() function at all taxonomic levels in *phyloseq*. Sample counts were then transformed to relative abundances. A filter was applied to subset phylogenetic groups in abundance greater than 2%. Each core microbiome was determined using the *microbiome* package in R (Lahti, Shetty, & Blake, 2017). The core() function in the *microbiome* package (detection = 0, prevalence = 0.9 and 0.1, respectively) was applied at each taxonomic level to determine the core taxa among all genotypes.

### Heritability Estimates

Heritability is an estimate that describes the proportion of phenotypic variance that is due to genetic variance. Narrow-sense SNP-based heritability (*h*^2^) was estimated using the *sommer* package in R as: 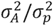, where 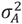 is the additive variance and 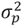 is the total phenotypic variance (additive + dominance) (Covarrubias-Pazaran, 2016). The core function of the *sommer* package is the ‘mmer’ function that fits multivariate linear mixed models. The modified exome capture SNP dataset, described above, was used to generate a kinship matrix used in ‘mmer’ to estimate narrow-sense heritability for each of bacterial families’ abundance. The *sommer* package includes a tolerance parameter for the matrix inverse when singularities are encountered in the estimation procedure. Here, inversion tolerance parameters were adjusted to 10 to avoid model singularity.

### Redundancy Analysis (RDA) and variance partitioning

To determine how whole microbiome variation changed across host genomic variation, much of which is due to population structure, we implemented redundancy analysis (RDA). RDA models variation in a set of response variables as a function of a set of explanatory variables. We used the first 10 principal components of switchgrass SNPs (Evans et al. 2014) as explanatory variables that describe population structure in switchgrass (Brown, Bray, & Pachter, 2018) and log-scaled ASV counts for each switchgrass genotype as the response variables. We performed the RDA to explain the relative proportion of total genotypic variance on the overall microbiome composition. The rda() function in vegan (Oksanen et al., 2019) was used to perform the RDA for the tetraploid accessions. For host genomic analyses like this RDA, we focused on tetraploids; since they (1) represent both upland and lowland ecotypes, (2) exhibit disomic inheritance unlike octoploids, and (3) exhibit more inter-group alpha diversity variation compared to octoploids.

We identified outlier ASVs with the strongest loadings, based on inter-quartile range criteria, on the first two RDA canonical axes; these ASVs are those most representative of microbiome-wide turnover across switchgrass hosts of different genomic background. ASVs that were labeled as outliers were then assigned taxonomic rank using the *phyloseq* method described above and compared to families in the original ASV dataset.

To dissect how genomic variation combined with ecotype, ploidy, and phenotypes to explain bacterial compositional turnover we also implemented variance partitioning of the RDA. Variance partitioning allows one to estimate the portions of compositional turnover explained by multiple sets of factors (Peres-Neto et al., 2006), which here were the first 10 principal components of SNPs, ecotype, geography-of-origin (latitude, longitude, and their squared values), and our two focal phenotypes (anthesis date and height). We implemented two versions of variance partitioning with RDA: one on ASV counts and the other on family abundances, to test whether family-level composition was differently associated with these factors than was ASV level composition.

### Genome-wide association study (GWAS)

To identify specific loci in the switchgrass genome linked to variation in rhizosphere bacterial composition, a genome-wide association study (GWAS) was conducted using the *statgenGWAS* package in R (Rossum et al., 2020). This fast single trait GWAS method was developed by Biometris, following the method described in Kang et. al (2010). The modified SNP dataset, described above, was once again used for this analysis.

Family-level abundance counts data (1) were imported from *phyloseq* using the rarified phyloseq object and a chromosome positional map (2) was generated from the SNP matrix (3). These three components were used to construct the R object used by *statgenGWAS*. Initial results from the heritability estimates and RDA indicated that certain core families of bacteria were influenced by switchgrass genotype. Therefore, to determine associated genes, a single trait GWAS, where the trait was the abundance of each of the families, was completed using a Generalized Least Squares (GLS) method for estimating the marker effects and corresponding p-values. Our GWAS included random effects correlated according to the kinship matrix, which was calculated with the VanRaden method (VanRaden, 2008). A p-value threshold is required for this method to minimize the False Discovery Rate (FDR) following the algorithm proposed by Brzyski et al. (2017). The method limits the number of SNPs in statistical linkage with each other passing FDR control. Significant SNPs were identified using a 0.01 threshold for the false-discovery rate (FDR). Linkage disequilibrium (LD) in switchgrass decays over kilobase length. Following Grabowski et al. (2017), we used a window of 25kb to identify genes potentially linked to significant SNPs. Thus, in this study, we defined SNPs within 25 kb as being “within LD of significant SNPs”. For each trait, *statgenGWAS* outputs significant SNPs and SNPs within the defined cutoff. A full description of the method can be found in Rossum et al. (2020).

### GO Enrichment Analysis

Significant SNPs associated with bacterial family abundance and positions within 25 kb of significant SNPs were then used to identify associated genes and gene ontologies. Here, we focused on three families: Xanthobacteraceae, Sphingomonadaceae, and Micromonosporaceae. The Xanthobacteraceae and Sphingomonadaceae families were chosen because they represent the high-fidelity core microbiome. Micromonosporaceae was chosen due to its high relative heritability within our panel and its importance in agriculture applications (Trujillo, Hong, & Genilloud, 2014). *Phytomine* was used to identify Gene IDs for each family using the *P. virgatum v1.1* reference genome at www.phytozome.net. Gene IDs for corresponding SNPs were then used to perform a Singular Enrichment Analysis (SEA) against the gene ID (ver. 4) background for *P. virgatum* using the agrigo v2 web program (Du, Zhou, Ling, Zhang, & Su, 2010). Default parameters were used.

## Results

### Amplicon Sequence Analysis

For 383 initial rhizosphere soil samples, 96,902 amplicon sequences variants (ASVs) were obtained following initial quality filtering and sequence processing in *dada2* (Callahan et al., n.d.). After sample pruning (removing non-bacteria from the taxonomy table) and rarefaction (1,000 sequences/sample) in *phyloseq*, 365 samples containing 31,181 ASVs remained. Counts for unique phylogenetic classifications are as follows: Domain: 1, Phylum: 26, Class: 59, Order: 142, Family: 268, Genus: 510.

### Alpha diversity analysis

We detected significant differences in alpha diversity (Chao1 and Shannon Diversity) between groups. Upland ecotypes, which are adapted to regions at higher latitudes and those geographically closer to our growing site, exhibited higher rhizosphere diversity compared to lowland ecotypes (Figure 1A, Upland-Lowland: *p.adj* (Shannon) = 1.58e^-2^). Tetraploids, which are present in both upland and lowland ecotype groups, also exhibited higher diversity overall, compared to octoploids, which are exclusively upland (Figure 1B, 4X-8X: *p.adj* (Shannon) = <0.001). Finally, the North genetic cluster, primarily comprised of upland tetraploids, exhibited the highest rhizosphere diversity among all of the genetic clusters and there was a significant difference between the East and West genetic clusters (Figure 1C, North-East: *p.adj* (Shannon) = <0.001, North-West: *p.adj* (Shannon) = <0.001, North-Northeast: *p.adj* (Shannon) = <0.001, North-South: *p.adj* (Shannon) = <0.001, East-West: *p.adj* (Shannon) = 5.92e^-2^). All diversity measurements for accessions can be found in Supplemental Table 1.

**Figure 1:**
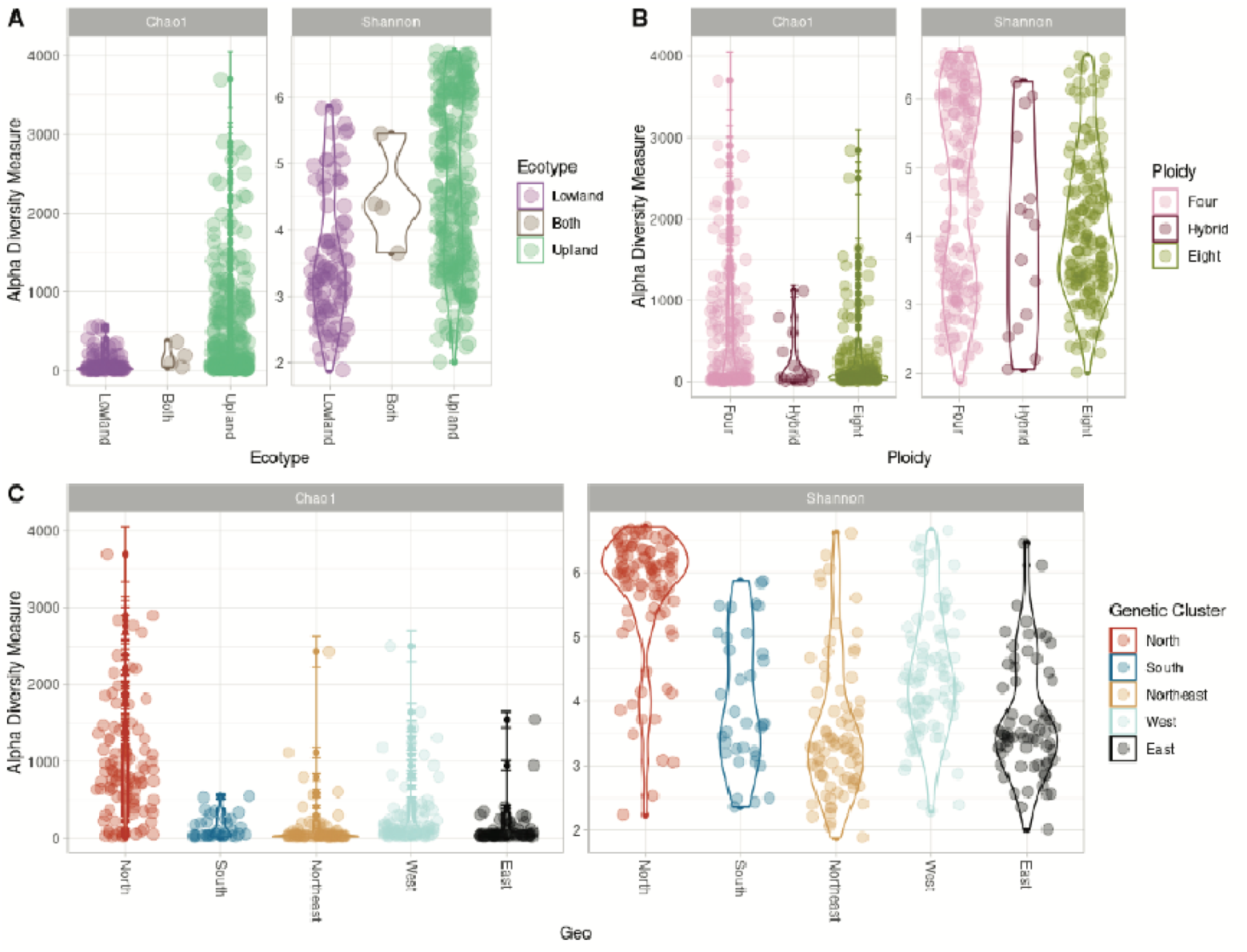
**A: Alpha diversity by Ecotype.** Alpha-diversity, measured by Chao1 and Shannon diversity Index, plotted for switchgrass ecotypes with Lowland (purple), Both (brown), and Upland (green). ***B: Alpha diversity by Ploidy.*** Alpha-diversity, measured by Chao1 and Shannon diversity Index, plotted for switchgrass ploidy levels with Tetraploid - Four (pink), Hybrid (maroon), and Octoploid - Eight (olive). ***C: Alpha diversity by Genetic cluster.*** Alpha-diversity, measured by Chao1 and Shannon diversity Index, plotted for switchgrass genetic clusters with North (red), South (dark blue), Northeast (beige), West (light blue), and East (black).

We found that Chao1 and Shannon diversity were negatively correlated with anthesis date (Chao1: *r* = - 0.17, p-value = 0.0014; Shannon: *r* = -0.18, p-value = <0.001) and full plant height (Chao1: *r* = -0.18, p-value = 0.0012; Shannon: *r* = -0.13, p-value = 0.017). We also considered these trait-diversity associations stratified by ecotype, ploidy, and genetic cluster (Figure 2). In general, the negative diversity association with anthesis date and height was broadly consistent, with Shannon diversity negatively correlated with plant height for Upland ecotypes (*r* = -0.25, p-value = <0.001), octoploids (*r* = -0.23, p-value = 0.0045), hybrids (*r* = -0.73, p-value = 0.016), the North genetic cluster (*r* = -0.27, p-value = 0.011), the Northeast genetic cluster (*r* = -0.22, p-value = 0.061), and the West genetic cluster (*r* = -0.24, p-value = 0.032) (Figure 2A,B,C). Likewise, Shannon diversity negatively correlated with anthesis date for Upland ecotypes (*r* = -0.19, p-value = 0.002), tetraploids (*r* = -0.23, p-value = 0.0025), hybrids (*r* = -0.94, p-value = <0.001), and the Northeast genetic cluster (*r* = -0.45, p-value = <0.001) (Figure 2D,E,F). A subset (n = 7) of ecotype/ploidy/genetic cluster combinations showed positive trait correlations with Shannon diversity, but these were not significant.

**Figure 2:**
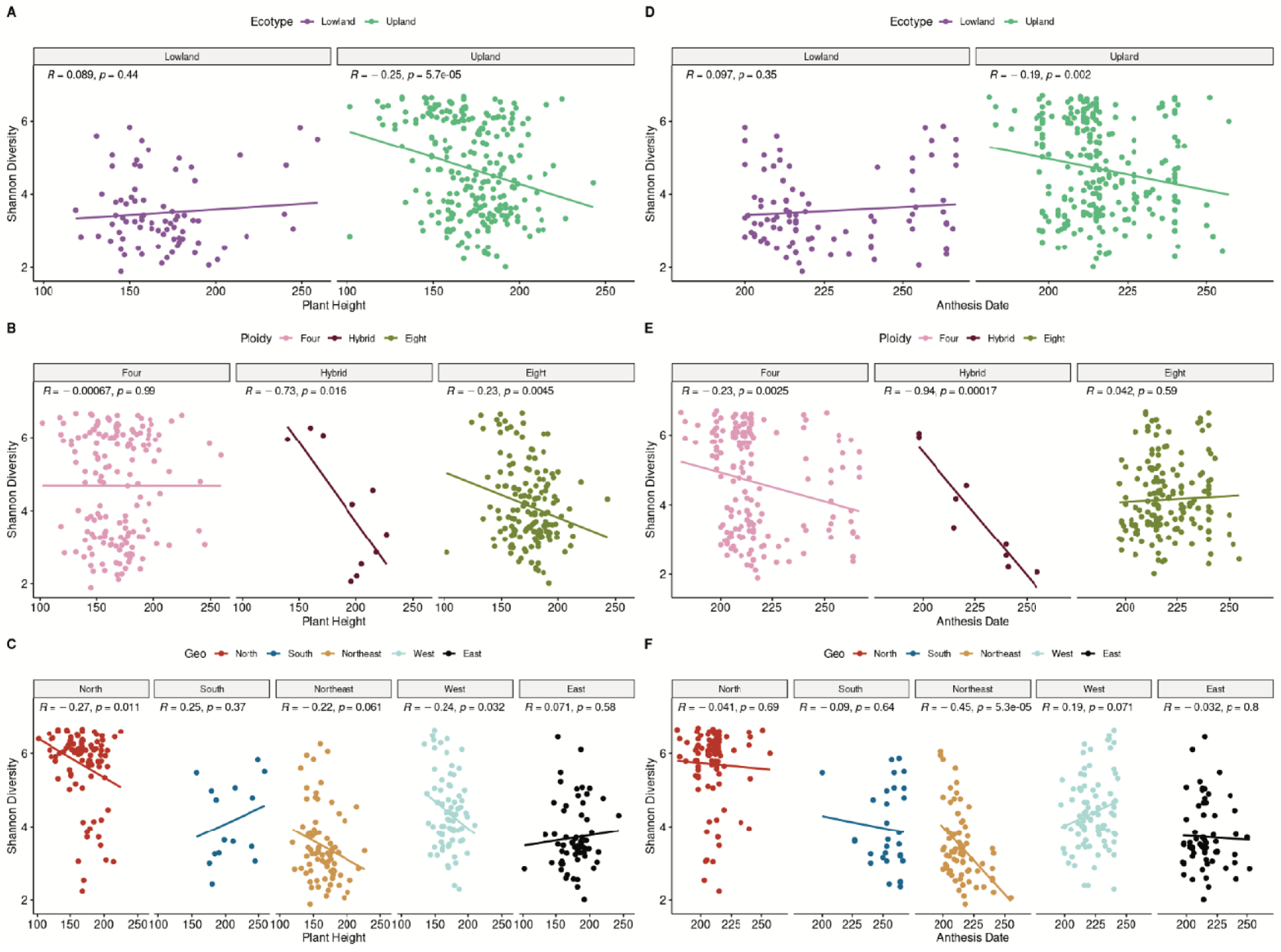
**Relationships between Shannon diversity and plant height. A: Ecotype**: plotted for switchgrass ecotypes wit Lowland (purple), Both (brown), and Upland (green). ***B: Ploidy:*** plotted for switchgrass ploidy levels with Tetraploid - Four (pink), Hybrid (maroon), and Octoploid - Eight (olive). ***C: Genetic cluster.*** plotted for switchgrass genetic clusters with North (red), South (dark blue), Northeast (beige), West (light blue), and East (black). ***Relationships between Shannon diversity and anthesis date. D: Ecotype***: plotted for switchgrass ecotypes wit Lowland (purple), Both (brown), and Upland (green). ***E: Ploidy:*** plotted for switchgrass ploidy levels with Tetraploid - Four (pink), Hybrid (maroon), and Octoploid - Eight (olive). ***F: Genetic cluster:*** plotted for switchgrass genetic clusters with North (red), South (dark blue), Northeast (beige), West (light blue), and East (black).

### Beta diversity analysis

We performed separate multivariate analyses of variance on a Bray-Curtis distance matrix using the NPMANOVA test. There were significant differences in overall bacterial composition between ecotypes (*p* < 0.001), ploidy levels (*p* < 0.001), and genetic clusters (*p* < 0.001), indicating differences in overall composition associated with switchgrass genetics and life-history.

Although all taxonomic levels were tested, the family level was identified as the lowest taxonomic rank with the ability to retain a high-fidelity core (Supplemental Figure 2). The high-fidelity core microbiome (>=90% of samples) contained two families: Xanthobacteraceae and Sphingomonadaceae. The low-fidelity core microbiome (>=10% of samples) contained 110 families (Supplemental Table 2).

### Host Genomic Influence

Of the 110 families in the core microbiome, twenty-one showed narrow-sense SNP-based heritability (*h^2^)* significantly greater than zero (Figure 3). The standard errors of narrow-sense heritability estimates are very large and there is considerable uncertainty in the estimate overall (Furlotte, Heckerman, & Lippert, 2014). Therefore, we consider estimates with standard errors that do not intersect zero as showing evidence of heritability (i.e. heritable). Heritability estimates for low-fidelity core taxa in the switchgrass rhizosphere ranged from 0.106 to 0.539, with a mean of 0.241. Heritability estimates presented here for bacterial lineages have a range similar to that of corn and sorghum (Deng et al., 2021). The heritability estimates indicate that, in part, switchgrass genotype influences the variability in the relative abundance of certain families within our growing site. Heritability estimates for all bacterial families can be found in Supplemental Table 2.

**Figure 3:**
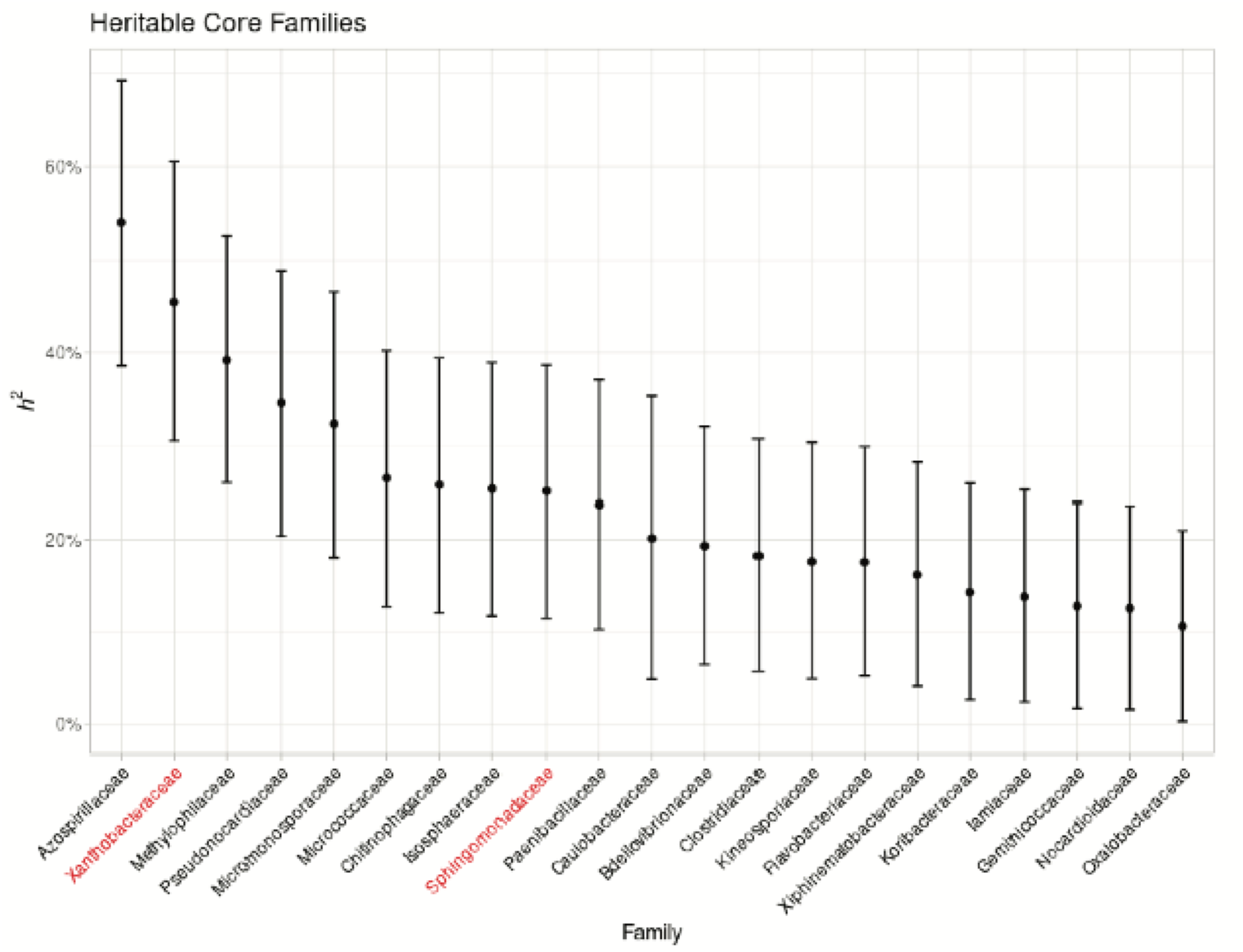
Heritability estimates for bacterial families in the switchgrass rhizosphere. For 110 bacterial families that were observed in more than 10% of switchgrass samples, narrow-sense heritability (*h^2^*) was calculated using a mixe linear model. 21 families had heritability estimates with standard error greater than zero. Families observed in more than 90% of our samples are represented in red.

Next, we performed a redundancy analysis (RDA) to characterize how the whole rhizosphere bacterial assemblage changes in association with host genomic background (Figure 4, Supplemental Figure 3). The RDA results indicate that genotypic variability among tetraploids (i.e. first 10 PCs of SNPs) had modest influences on bacterial ASV composition (*R^2^* = 0.063, *R^2^*_adjusted_= 0.007). Switchgrass SNP PC1, which is associated with separation between the North and the Northeast and South genetic clusters, loaded most strongly to the first primary canonical axis (RDA1, PC1 = 0.99, the next strongest loaded PC was PC10, with a 0.097 loading). Switchgrass SNP PC7, which is associated the variance in the North genetic cluster, loaded most strongly to the second canonical axis (RDA2, PC7 = 0.899). The proportion of explained variance in bacterial ASV composition was fairly low along these individual canonical axes (RDA1 = 0.9%, RDA2 = 0.7%), which was unsurprising given the large diversity in the bacterial composition. Still, the RDA identified bacterial ASV compositional turnover in association with switchgrass population structure, as shown by the turnover between the North and the Northeast and South switchgrass genetic clusters (Figure 4).

**Figure 4:**
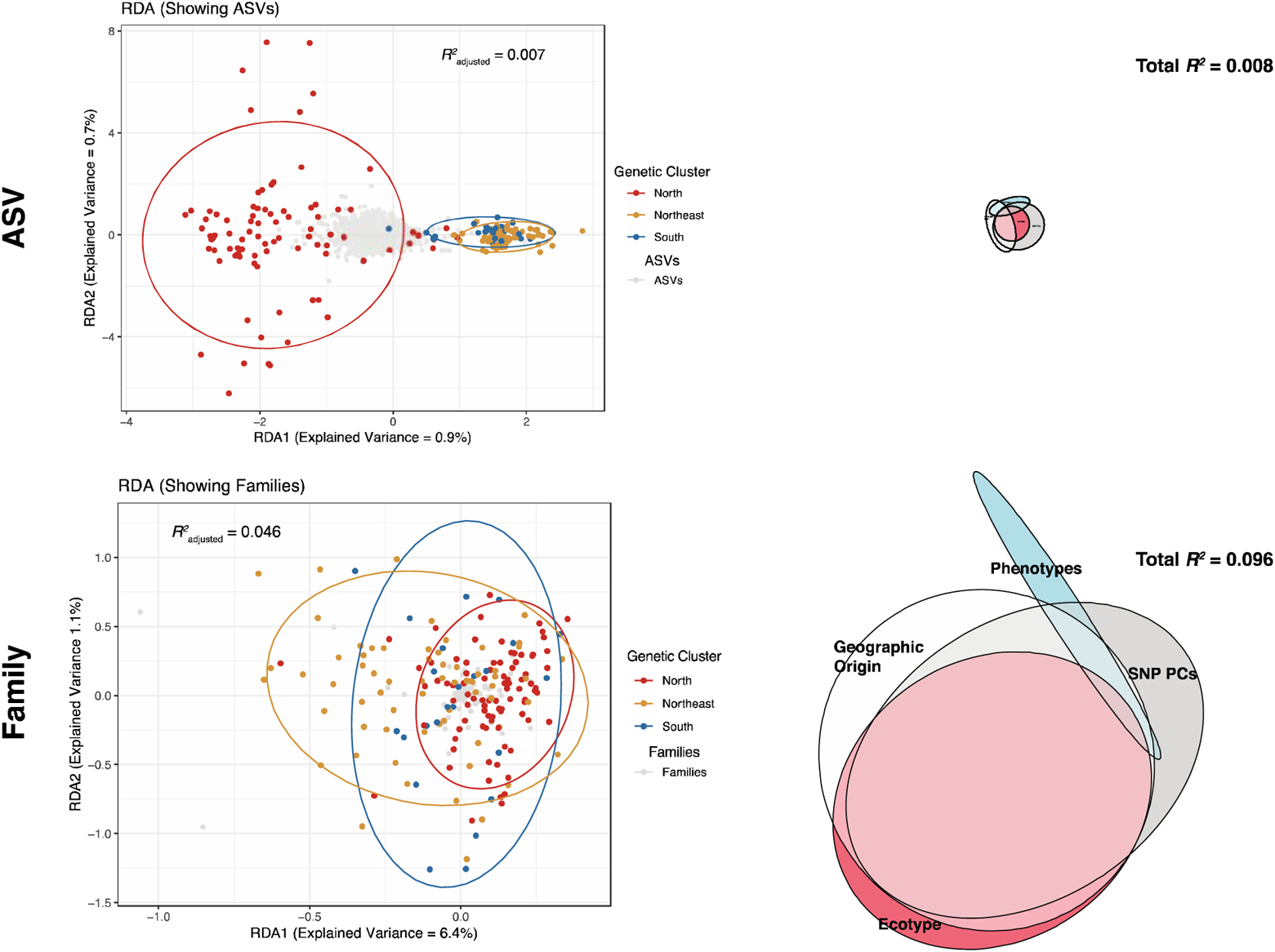
**Redundancy Analyses and Variance Partitioning. *A:*** Results of RDA analysis presenting amplicon sequence variants (Grey) and the North (Red), Northeast (Beige), and South (Blue) genetic clusters. ***B:*** Results of variance partitioning analysis for ASVs presenting Ecotype (Red), SNP PCs (Grey), Phenotypes (Blue), and Geographic Origin (White). ***C:*** Results of RDA analysis presenting rhizosphere bacterial families (Grey) and the North (Red), Northeast (Beige), and South (Blue) genetic clusters. ***D:*** Results of variance partitioning analysis for rhizosphere bacterial families presenting Ecotype (Red), SNP PCs (Grey), Phenotypes (Blue), and Geographic Origin (White).

**Figure 5:**
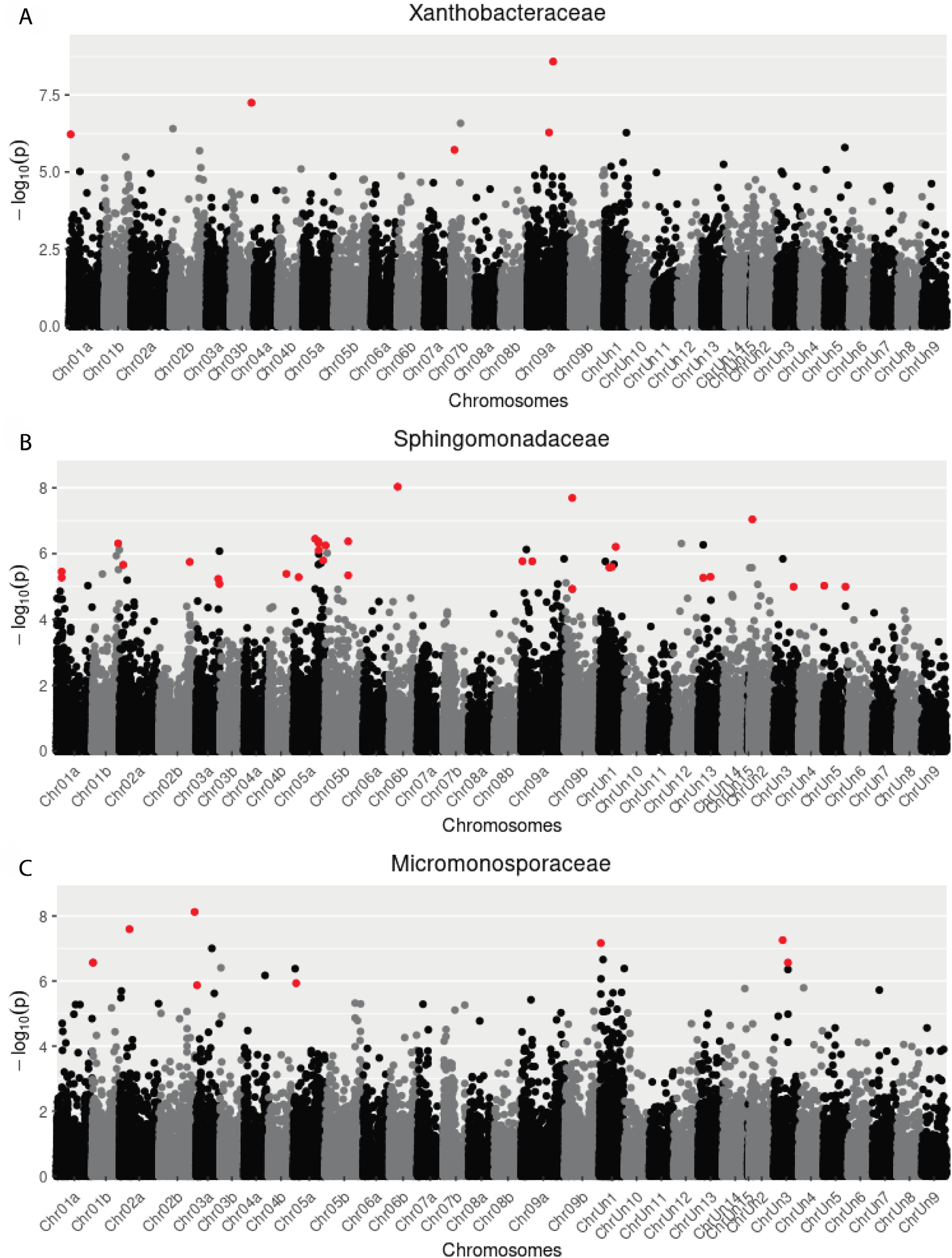
Genome-wide association study (GWAS): A: Manhattan plot presenting significant SNPs (Red) associated with Xanthobacteraceae rhizosphere abundance. ***B:*** Manhattan plot presenting significant SNPs (Red) associated with Sphingomonadaceae rhizosphere abundance. ***C:*** Manhattan plot presenting significant SNPs (Red) associated with Micromonosporaceae rhizosphere abundance.

Outlier ASVs were determined for each of the first two canonical RDA axes (n = 9,215). These outlier ASVs are most representative of the variation in bacterial composition among the tetraploid genotypes. We then identified the taxonomic ranks for the outlier ASVs. Our results show that Xanthobacteraceae and Sphingomonadaceae represent the high-fidelity core for both the outlier ASV dataset and the original ASV dataset, signifying host genotypic influence might be strongest on these families. When we consider the low-fidelity core (n = 110 families), 70.9% of families in the RDA outlier ASV core microbiome are in original core microbiome (Supplemental Table 2). When we consider just the heritable families, 80.95% of heritable families in the outlier ASV core microbiome are in the original core microbiome (Supplemental Table 2).

When we applied variance partitioning to estimate the portion of ASV-level compositional turnover explained by SNP PCs, ecotype, geographic origin (i.e. latitude and longitude coordinates), and the two phenotypes, we found that the total *R^2^* was low, approximately %1 of ASV composition. However, variance partitioning of family-level turnover found these 4 factors explained a larger portion, with *R^2^* = 0.096 (Figure 4). This variance partitioning found that the largest portion of bacterial compositional variation was explained by the collinear portion of SNP PCs, ecotype, and geographic origin (R^2^ = 0.0866), suggesting that geographic population structure associated with ecotypic variation explains a large portion of turnover in bacterial family composition.

### Genome-wide association study

A primary limitation of heritability estimates and the RDA was that we could not account for which SNPs have the greatest effect on compositional variation. To overcome this, a genome-wide association study (GWAS) was conducted to identify specific loci associated with the relative abundance of families in the core microbiome. In total, 1,861 SNPs were found to be associated with the abundance of the 110 tested core families. 878 SNPs were found to be associated with the abundance of the 21 heritable core families. Furthermore, 36 SNPs were found to be associated with the abundance of the two families (Xanthobacteraceae and Sphingomonadaceae) that represent the high-fidelity core microbiome (Figure 6, Supplemental Table 2).

### GO Enrichment Analysis

Despite the general difficulty identifying specific molecular functions or pathways in non-model plant species, resources for gene ontology (GO) still remain useful for biological inference (Tian et al., 2017). Therefore, we completed a GO enrichment analysis for genes harboring significant SNPs and positions within 25kb of significant SNPs to determine possible cellular processes (P), cellular components (C), and molecular functions (F) associated with three core family abundances (Supplemental Table 3). Primarily, we identified two types of cellular processes: metabolic and stimuli response processes. Broadly, we identified several molecular functions relating to ATPase activity, hydrolase activity, and pyrophosphatase activity (Supplemental Figure 4, Supplemental Table 3). Functions involving ATP hydrolysis as an energy source either typically catalyze a reaction or drive membrane transport against a concentration gradient (Berg, Tymoczko, & Stryer, 2002). In the context of our GWAS, this could suggest possible root exudate activity whereby switchgrass genotypes differentially transport exudates against a lower soil compound gradient influencing core family abundances. Perturbing these regions in future studies could reveal the underlying pathways and mechanisms. Lastly, genes harboring SNPs associated with Sphingomonadaceae abundance, in particular, were enriched for GO terms associated with cellular membrane components, metallopeptidase and metalloendopeptidase activity, and cellular processes that relate to metabolism and response to stimuli (Supplemental Table 3). Again, given the context of our GWAS, we suspect that the genes associated with these GO terms could interact with Sphingomonadaceae populations in the soil. More work involving these genes could expose perturbations that more predictably affect Sphingomonadaceae abundance in switchgrass.

## Discussion

Macroscopic organisms often exhibit striking variation among individuals in the microbial assemblages they host, but the genetic basis and ecological relevance of this variation is often unclear. Common garden experiments with diverse genotypes offer a powerful window into dissecting the ecological genomics of host influences on their microbiomes. Previous studies have demonstrated that genomic differences among switchgrass genotypes and populations may, in part, have led to much of the adaptive phenotypic differences observed between switchgrass populations (Adkins et al., 2016; Lowry et al., 2019). We add to these observations by showing a similar phenomenon within a less visible portion of the switchgrass phenotype. We found that switchgrass genotype influences the rhizosphere microbiome in a variety of ways relative to bacterial alpha and beta diversity as a result of variability within the switchgrass genome.

We demonstrated that the alpha diversity (Chao1 and Shannon diversity) of rhizosphere bacteria within a common garden was significantly differentiated between switchgrass ecotypes, ploidy levels, and genetic clusters. These groupings with distinct bacterial assemblages are associated with life history variation and locally-adapted traits in this species. Furthermore, alpha diversity within each group typically had no correlation with, or was negatively correlated with, anthesis date and plant height (Figure 2). Flowering time is an important adaptation to local growing conditions and has major fitness consequences for the plant (Grabowski et al., 2017b). In general, in common garden experiments, upland ecotypes have an anthesis date 2–4 weeks earlier than lowland ecotypes (Schwartz & Amasino, 2013). As plants move through different developmental stages the diversity and availability of metabolites influencing the rhizosphere microbiome shifts (Wagner, 2021). Rhizosphere sampling in this study occurred in May, 2016. Therefore, it’s reasonable to expect genotypes at different stages of development at the time of sampling to exert different selective pressure on the microbiome composition. Future research examining temporal exudate availability in the rhizosphere of switchgrass could expand on this hypothesis.

It is already widely hypothesized that hosts mediate their microbiomes relative to environmental conditions, resource availability, and microbial function, not necessarily always a specific species or strain (Lemanceau et al., 2017; Martiny et al., 2015). Thus, some basis for comparing taxa between host groupings becomes important when determining the taxonomic level at which host influence is detectable. A “core” microbiome can be defined in a variety of ways, and may not represent microbes that have the most important functional impacts on a system (Bell & Bell, 2021). For example, commensal microorganisms remain widespread throughout host populations (Zeng et al., 2015). Still, the consistent occurrence of microbial taxa provides an opportunity to assess relationships between host genetics and microbial recruitment. Here, we applied the “common core” concept (Risely, 2020) to identify bacterial taxa that were consistently associated with diverse switchgrass genotypes in a common garden. Given the disparity in evolutionary pace (i.e. there are many bacterial generations during the lifecycle of a switchgrass plant), it is unlikely that switchgrass has mechanisms to distinguish between closely related bacterial taxa unless doing so provided a substantial fitness advantage (Yin et al., 2021). Therefore, the core microbiome provides both a basis for measuring bacterial abundance across multiple switchgrass genotypes as well as a means to detect host genomic influence at the lowest possible taxonomic rank.

For example, Singer et al. (2019) observed a large core endophytic switchgrass microbiome dominated by root-colonizing bacterial genera such as *Streptomyces*, *Pseudomonas*, and *Bradyrhizobium*, while rhizosphere diversity was more variable between their two sampling sites. Likewise, Singer et al. (2019b) observed different core bacterial classes associated with upland and lowland switchgrass ecotypes, with each ecotype preferentially enriched for Alphaproteobacteria and Actinobacteria, respectively. For our study, out of 268 families, only two associated with the majority (>=90%) of our rhizosphere samples (Xanthobacteraceae and Sphingomonadaceae) and less than half (n = 110) associated with more than 10% of our samples. Xanthobacteraceae and Sphingomonadaceae are families in the Alphaproteobacteria class. We were unable to detect any genera or species associated with our genotypes at the 90% occupancy threshold. This indicates that a large swath of the switchgrass rhizosphere microbiome assembly is stochastic and only certain lineages were able to consistently associate with our genotypes in the common garden.

The abundance of twenty-one of the 110 families were determined to show evidence of heritability. After reasonably establishing heritability of select core families, the RDA allowed us to identify how genome-wide variation in switchgrass influenced the rhizosphere microbiome composition. RDA methods used in microbial ecology typically explain the variance in microbiome composition due to environmental variables, such as temperature or pH (Paliy & Shankar, 2016). However, here, we modeled microbial compositional data as a function of host genomic data. As such, the RDA in this study represents the compositional variance of the microbiome due to genetic variation in the hosts. We then extended the analysis to explore outlier ASVs along each of the first two RDA axes, because those ASVs represent taxa associated with host genomic influence on overall turnover in bacterial composition. Once clustered at the family level, we determined that seventeen out of the 21 heritable families were in the outlier ASV dataset. Additionally, the results of the variance partitioning suggest that geographic population structure associated with ecotypic variation explains a larger portion of turnover in bacterial family composition compared to ASV composition. Thus, we determined that genotypic variability in switchgrass sampled from across its natural range influences the composition of the rhizosphere bacterial microbiome, particularly for tetraploids.

The ecophysiology of host-microbe interactions in our study is unclear, requiring detailed investigation, but there may be hints in the function of switchgrass genes that are strongly associated with core family abundances. We conducted a GWAS to identify potential loci that strongly associate with core family abundances. The outcrossing nature of switchgrass and its rapid decay in LD can help pinpoint causal loci with GWAS. Several considerations had to be made though, in order to reliably trust the significance of the resulting QTLs. First, we considered only tetraploids for the same reasons as the RDA, that they (1) represent both upland and lowland ecotypes, (2) exhibit disomic inheritance unlike octoploids, increasing the effect size of additive effects, and (3) exhibit the most inter-group alpha diversity variation compared to octoploids or hybrids. We used exome capture SNPs (Evans et. al. (2018), which are more effective than genotype-by-sequencing (GBS) in their ability to tag causal polymorphisms, compared to GBS SNPs that are often far from any genes (Kaur et al., 2014). The identification of genes containing significant SNPs could provide insights into potential host-microbe interaction pathways. To explore this relationship further, we chose to focus on three core families: Xanthobacteraceae, Sphingomonadaceae, and Micromonosporaceae.

We identified hundreds of switchgrass genes containing SNPs (including those within 25 kb of significant SNPs) associated with Micromonosporaceae (n = 315), Xanthobacteraceae (n = 578), and Sphingomonadaceae (n = 718) abundance in the rhizosphere. Many species within the Micromonosporaceae family degrade chitin, cellulose, lignin, and pectin, and play an important role in the turnover of organic plant material. Moreover, many strains produce useful secondary metabolites and enzymes used by plants (Trujillo et al., 2014). Members of Xanthobacteraceae have been experimentally identified as nitrate reducers (C. R. Anderson et al., 2011) and therefore their regulation could be important to host metabolic activity. Our GO enrichment analysis indicates that genes associated with Micromonosporaceae and Xanthobacteraceae abundance are involved in hydrolase, pyrophosphatase, and ATPase activity suggesting that the enrichment for this set of genes plays an important role in overall plant fitness. Members of the Sphingomonadaceae family are known to be antagonistic against plant pathogens and induce plant growth promotion (Glaeser & Kämpfer, 2014). GO terms for genes associated with this family are enriched for metallopeptidase and metalloendopeptidase activity, membrane cellular components, and processes involved in stimuli response. Altogether, this suggests that the activity of these genes is linked to environmental stimuli, possibly initiated by Sphingomonadaceae, and interacting with plant cell membranes. Future work examining these families could make these connections to plant function more robust.

In conclusion, bacterial composition in the switchgrass rhizosphere is influenced by host genotypic variability. Rhizosphere diversity shows major differences among switchgrass ecotypes, ploidy levels, genetic clusters, and life-history axes. In particular, the North switchgrass genetic cluster was most distinct at our common garden site. Two high-fidelity core families, Xanthobacteraceae and Sphingomonadaceae, were observed across the majority of our switchgrass genotypes. Twenty-one bacterial families were found to exhibit evidence for heritability. Though the explained variance was low, a redundancy analysis revealed widespread genotypic influence on the composition of the rhizosphere bacterial microbiome and outlier ASVs along each primary axis belonged largely to our defined core microbiome. Finally, we identified 1,861 SNPs associated with the abundance of core microbiome families and discussed potential pathway mechanisms that could influence a subset of those families that we expected could influence host fitness. A GO enrichment analysis determined several functional and cellular processes for which host genotypic influence could be expressed, notably by hydrolase activity. This study should function as a foundation on which further exploration can be achieved investigating the relationship between switchgrass genotype and its associated rhizosphere microbiome in different environmental contexts or under artificial genetic perturbation.

## Acknowledgements

We thank Wanyan Wang for assistance in collecting and cataloging the soil samples from the GWAS panel and selecting samples for metagenome analysis. We thank Maureen Mailander for conducting the initial rhizosphere soil DNA extractions. We are also grateful to our entire project team (investigators Marvin Hall, Stacy Bonos, Julie Hanson, Don Viands, and their students and staff for the collection of the GWAS family plants from the common garden site at Ithaca, NY). We greatly appreciate the opportunity provided by Dr. Michael Casler for allowing us to collect samples of all of the genotypes in this range-wide collection of switchgrass genotypes and check cultivars at of Cornell University for this project. This research was supported by the National Institute of Food and Agriculture, U.S. Department of Agriculture, under award number 2019-67009-29006 to JEC, TBH, and JRL. Seed support was provided by the Northeast Woody/warm-season BIOmass consortium, funded by USDA-AFRI Grant #2012-68005-19703. Program support to JEC was provided through the USDA National Institute of Food and Agriculture Federal Appropriations under Project PEN04532 and Accession number 1000326. JRL was also supported by NIH award 1R35GM138300-01.

## Data Availability

Raw amplicon sequence reads are deposited in the SRA (BioProject PRJNA689762).

## Supplemental Figures

**Supplemental Figure 1:**
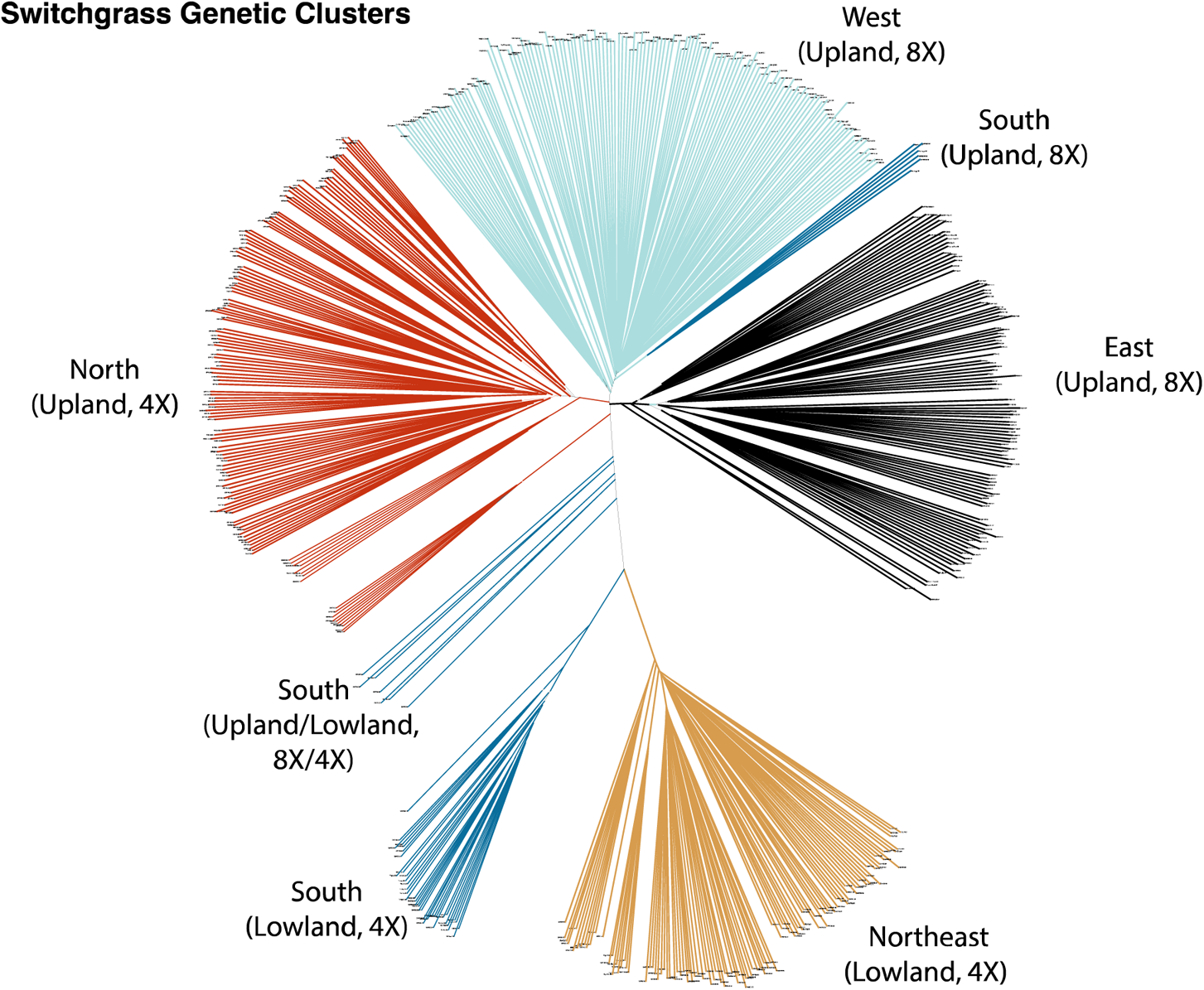
***Phylogenetic analysis:*** Results of phylogenetic analysis based on exome capture dataset from Evans et al. 2018 with switchgrass genetic clusters colored as North (red), South (dark blue), Northeast (beige), West (light blue), and East (black).

**Supplemental Figure 2:**
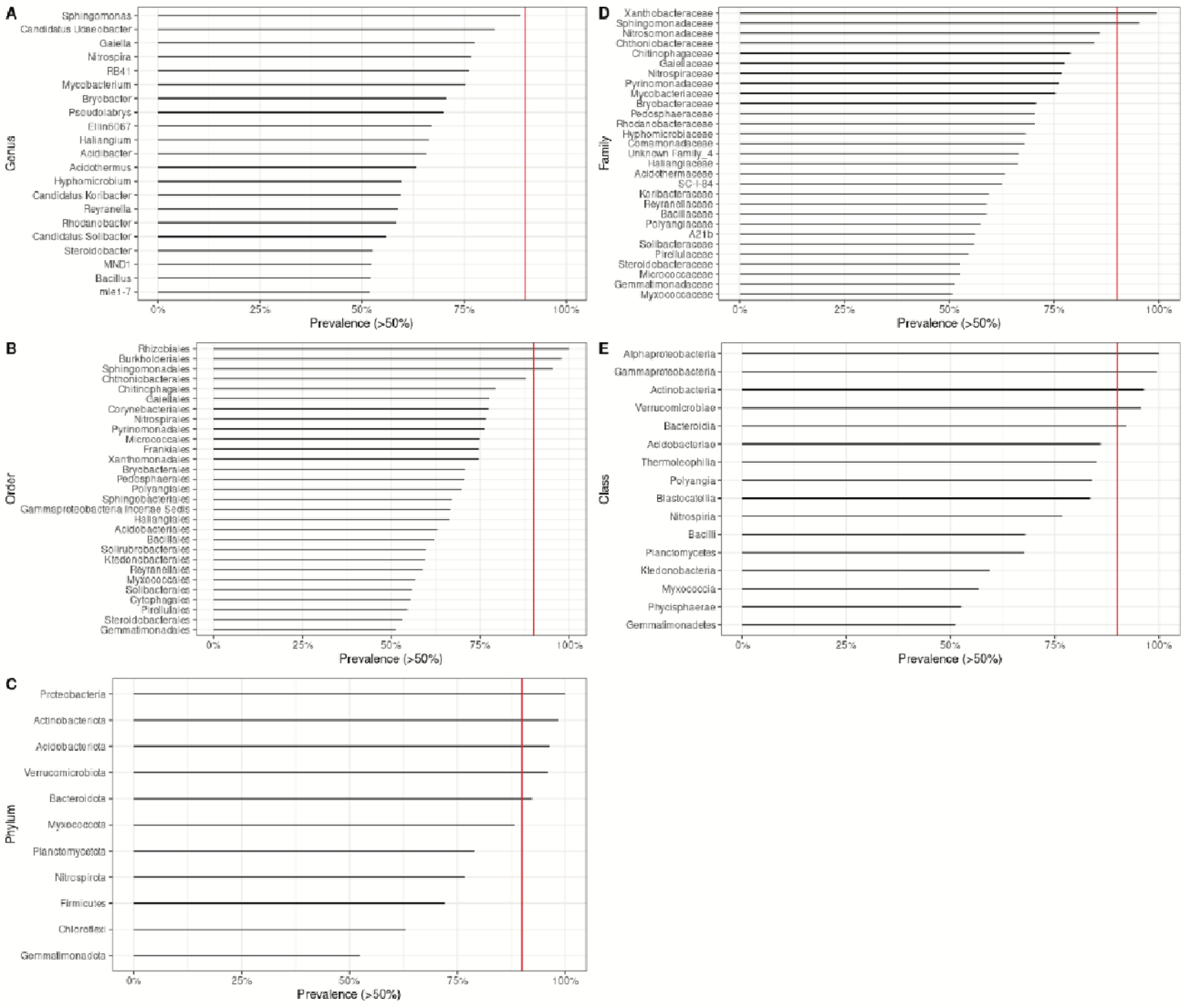
***Prevalence of rhizosphere bacterial taxa:*** The prevalence of rhizosphere bacterial taxa in greater than 50% of switchgrass samples shown for ***A:*** Genus, ***B:*** Family, ***C:*** Order, ***D:*** Class, and ***E:*** Phylum. The 90% occupanc threshold for each group is indicated by a vertical red line.

**Supplemental Figure 3:**
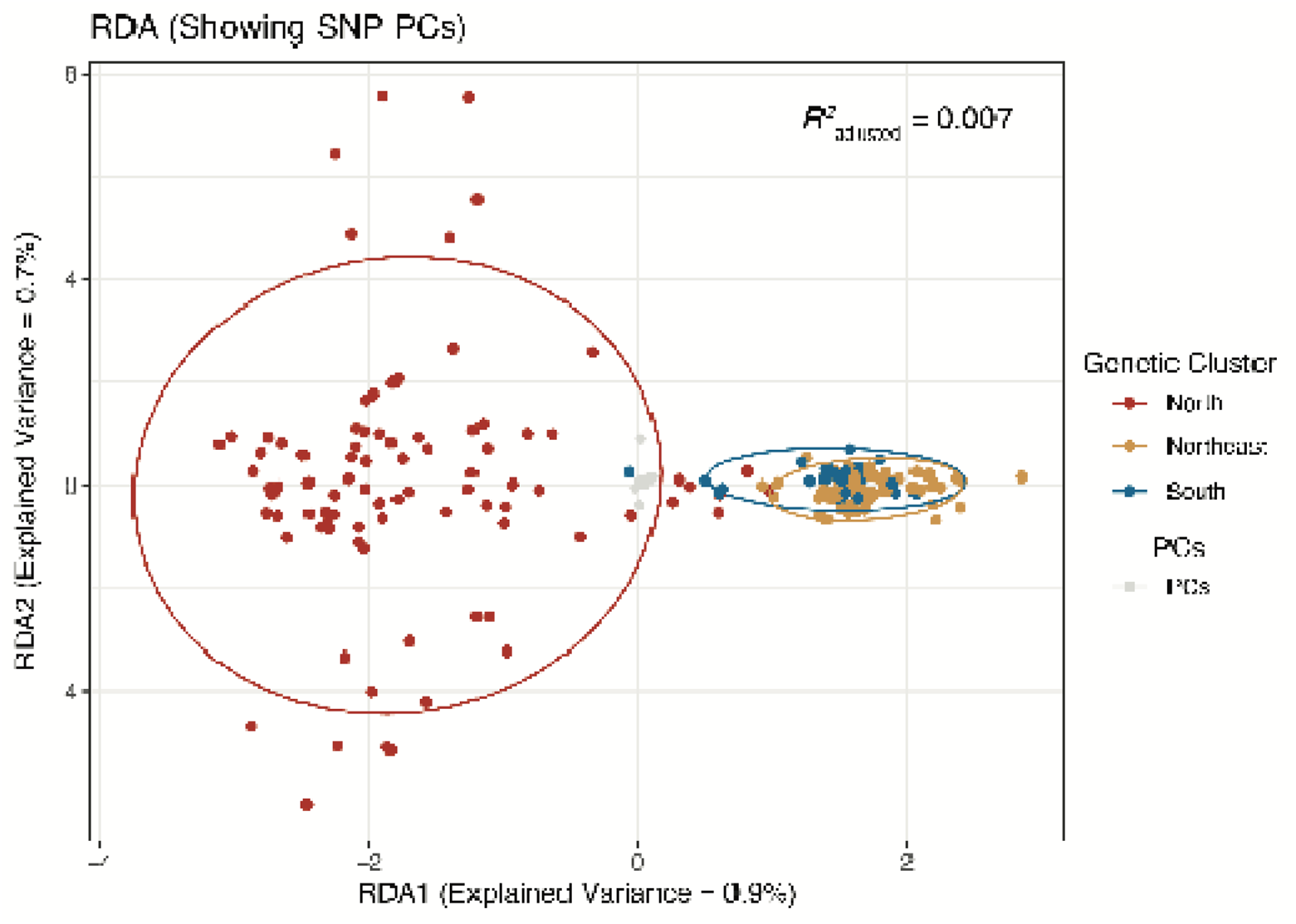
***Redundancy Analyses (PCs):*** Results of RDA analysis presenting SNP PCs (grey) and the North (red), Northeast (beige), and South (dark blue) genetic clusters.

**Supplemental Figure 4:**
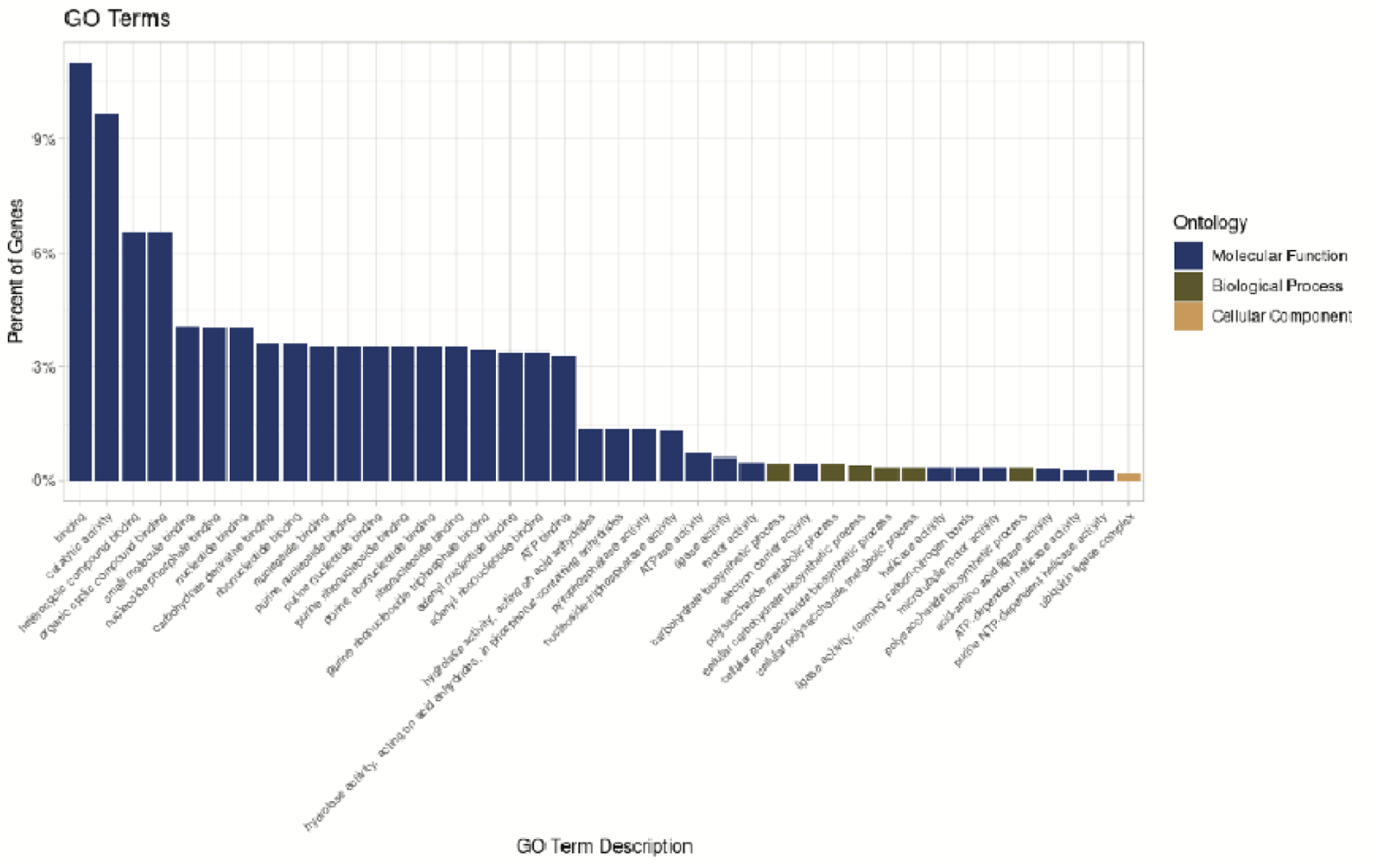
***GO Terms:*** Percent of enriched GO terms related to Molecular Function (dark blue), Biological Process (olive), an Cellular Component (beige).

**Supplemental Figure 5:**
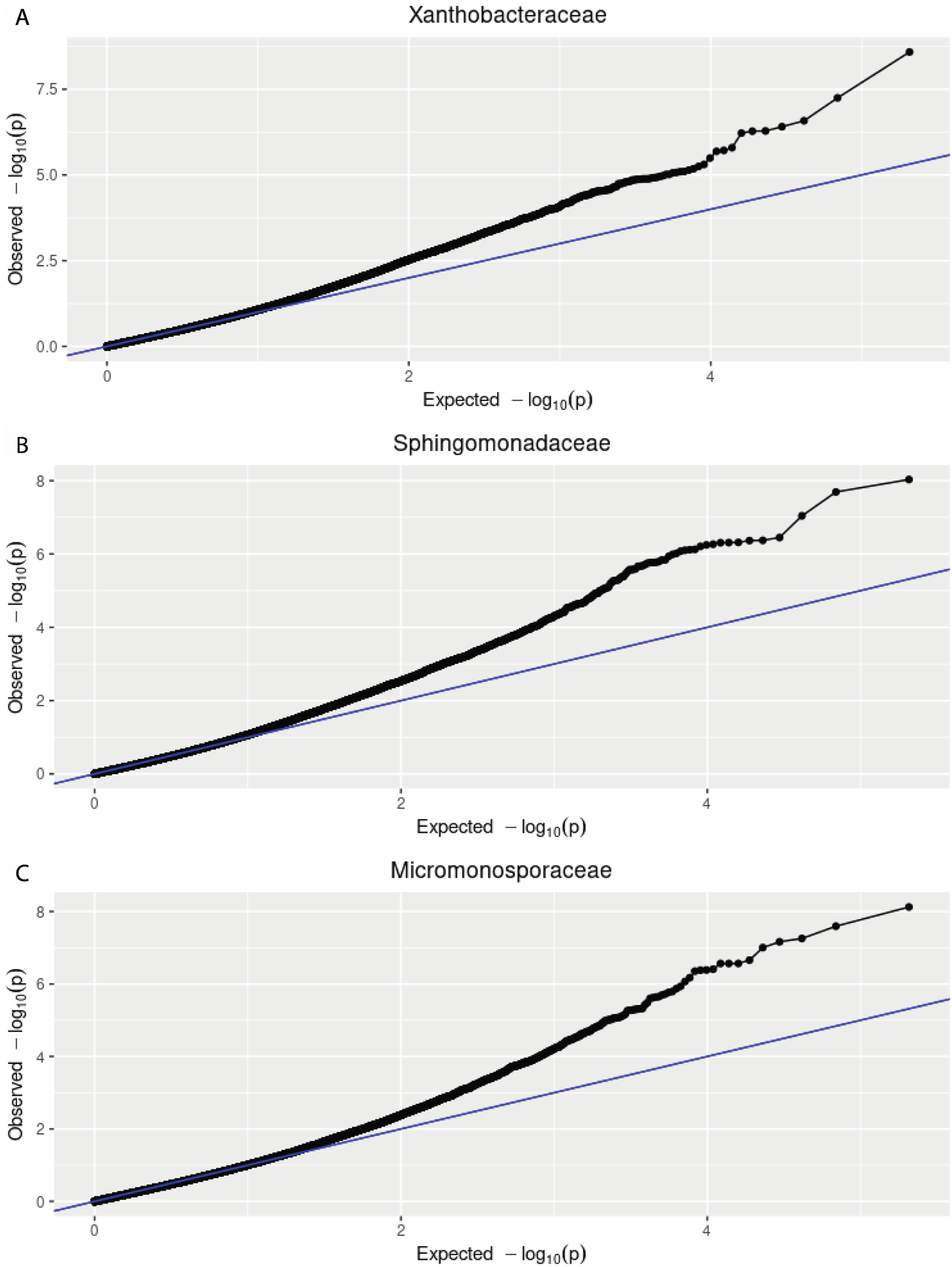
***Genome-wide association study (GWAS): A:*** QQ plot presenting significant SNPs (red) associated with Xanthobacteraceae rhizosphere abundance. ***B:*** QQ plot presenting significant SNPs (red) associated with Sphingomonadaceae rhizosphere abundance. ***C:*** QQ plot presenting significant SNPs (red) associated with Micromonosporaceae rhizosphere abundance.

## Supplemental Tables

**Supplemental Table 1:**
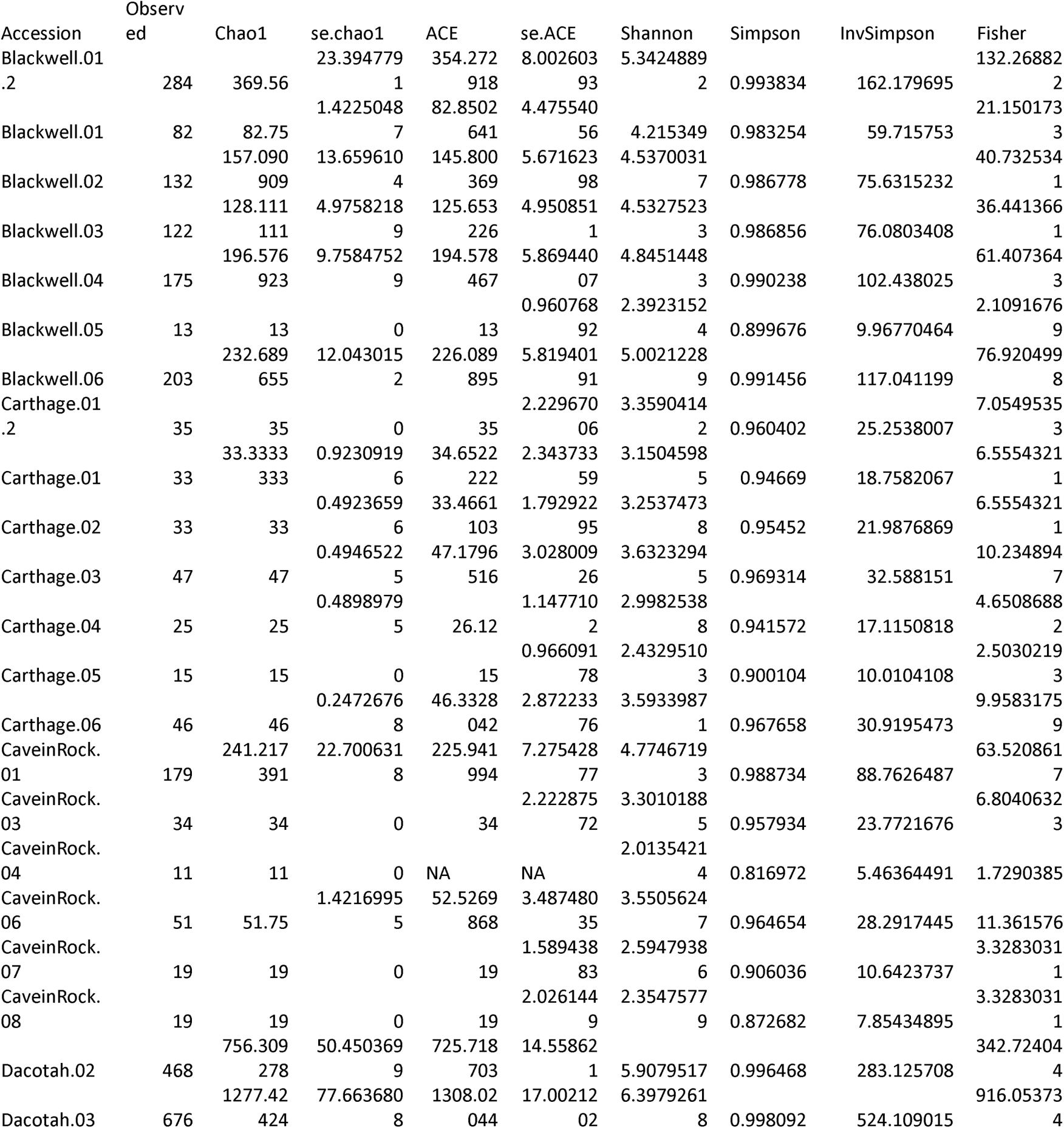
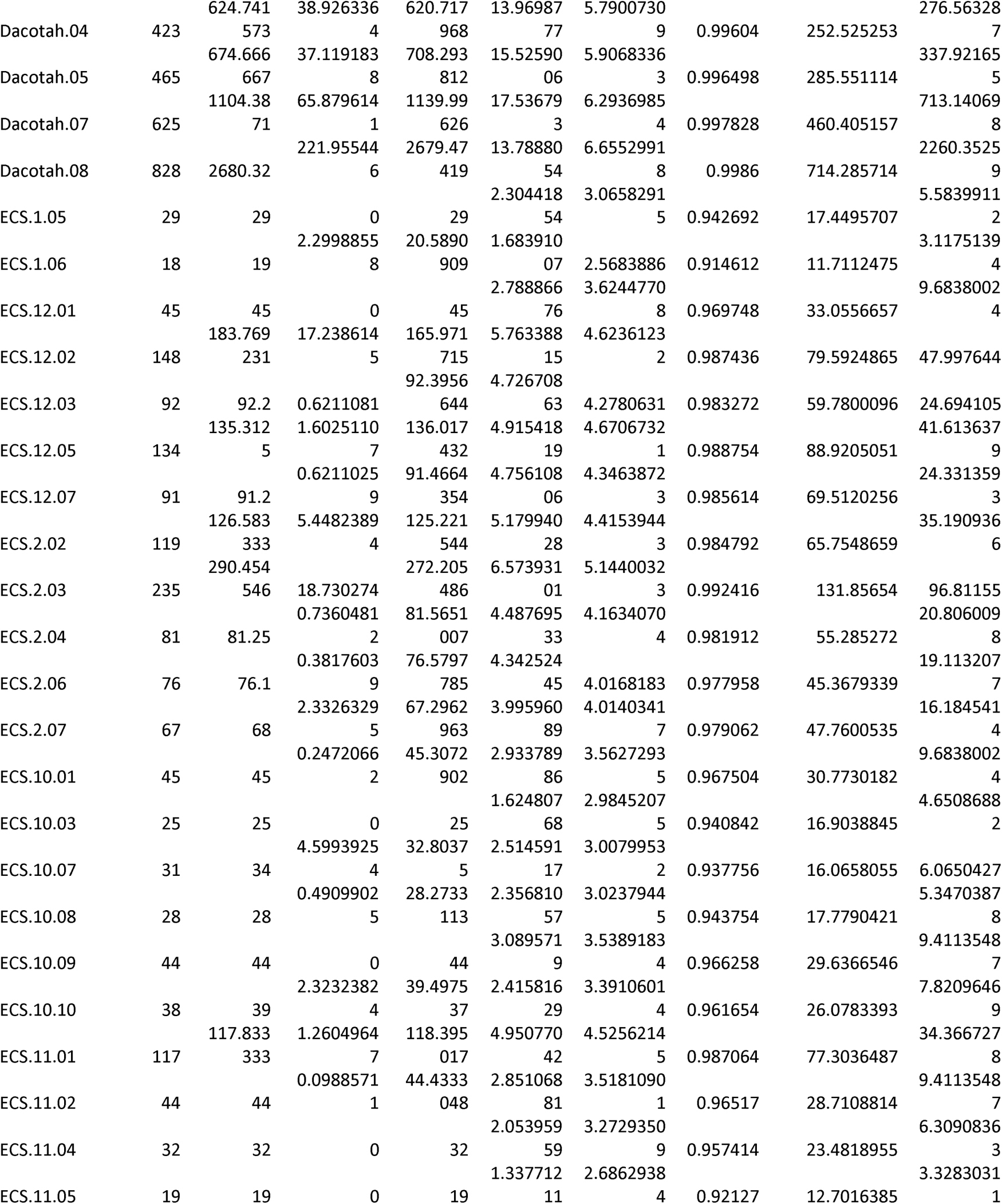
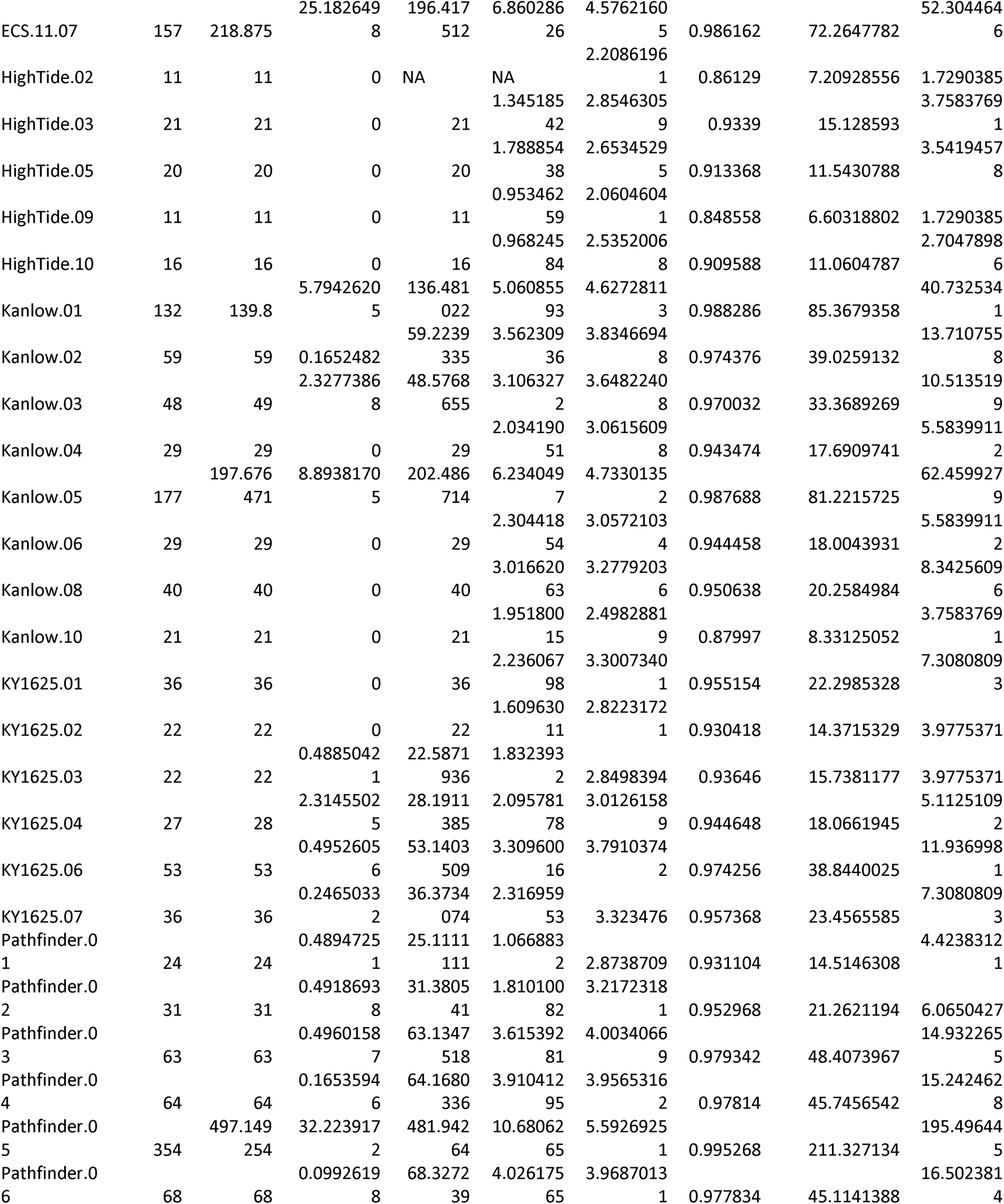
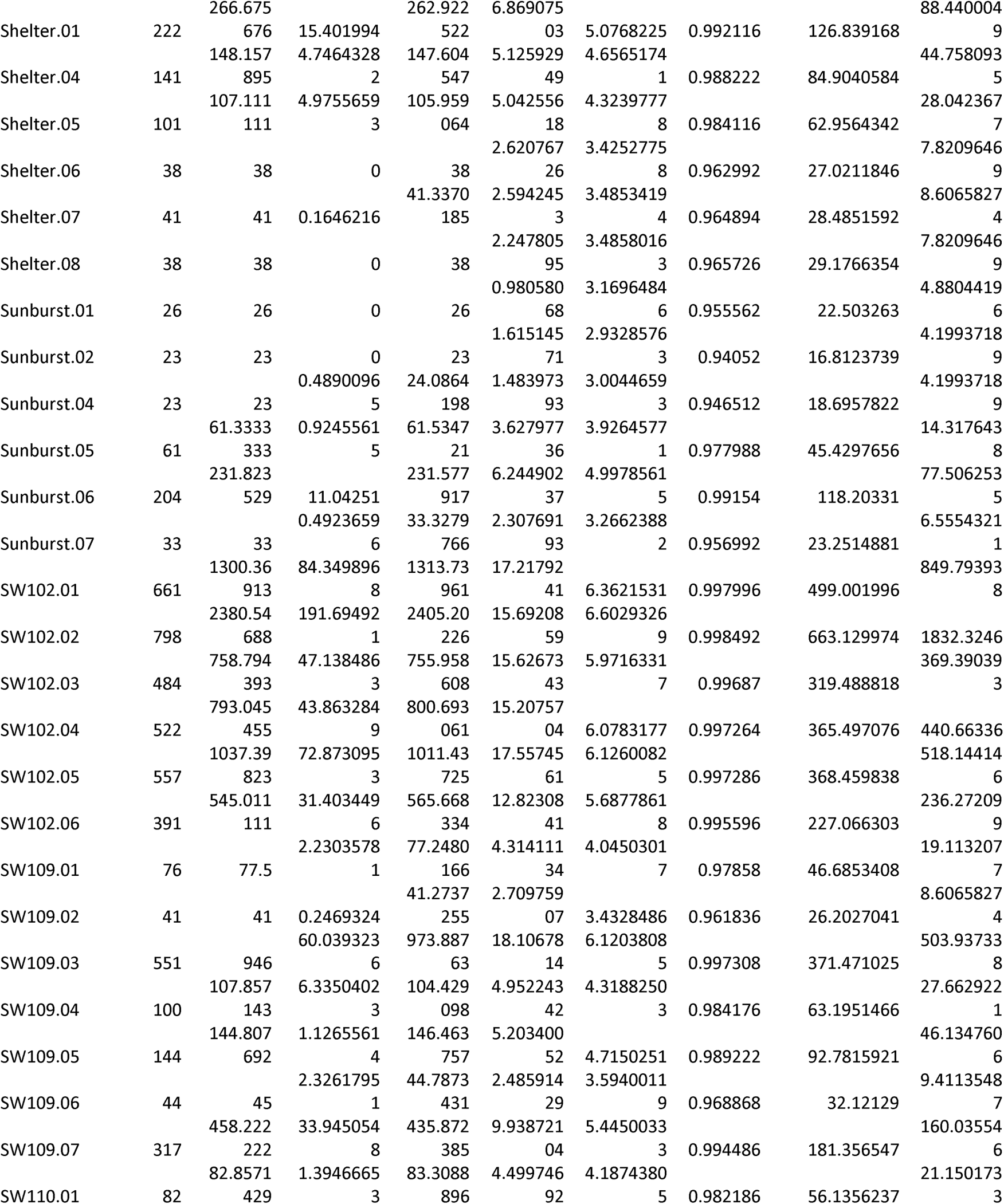
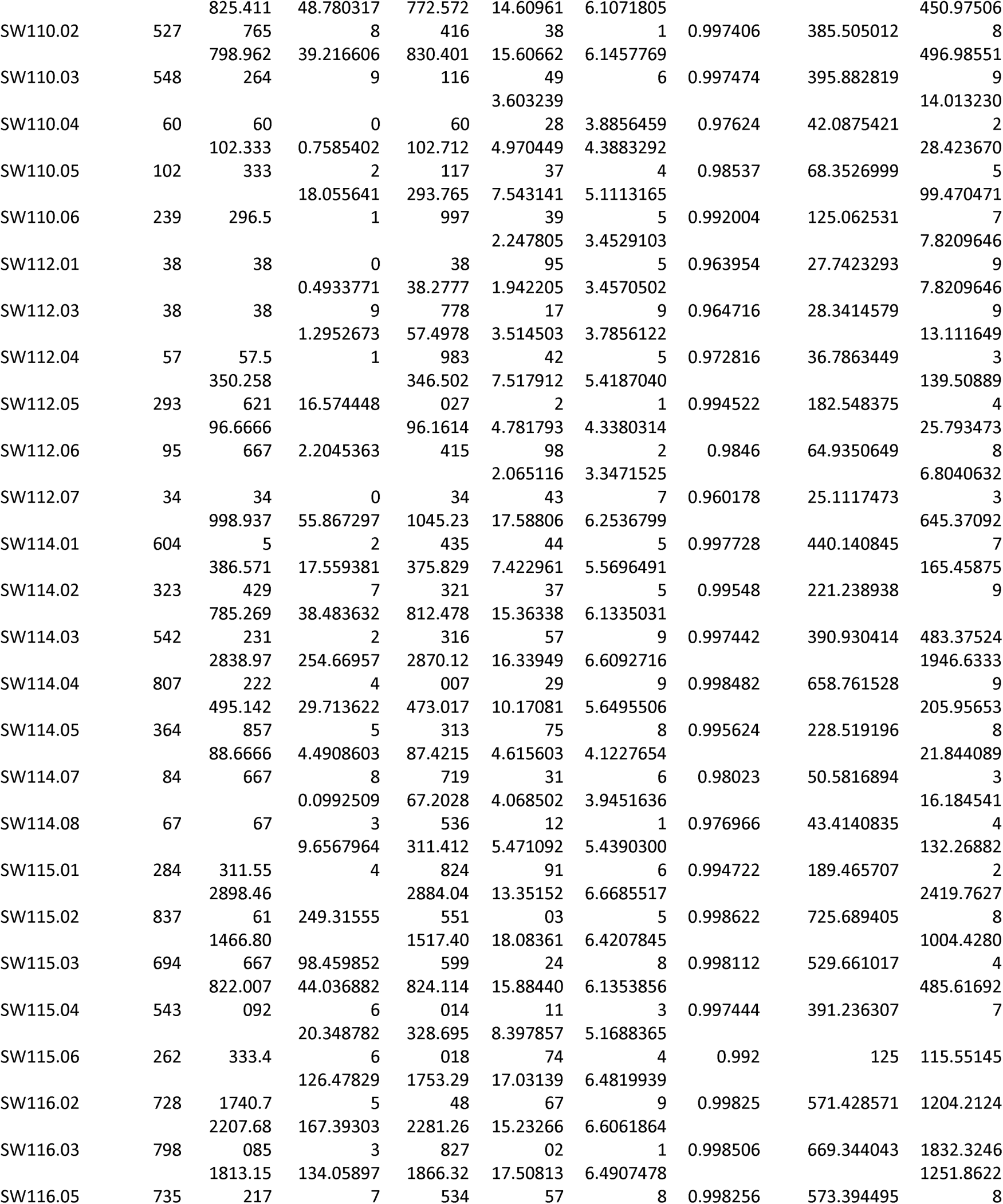
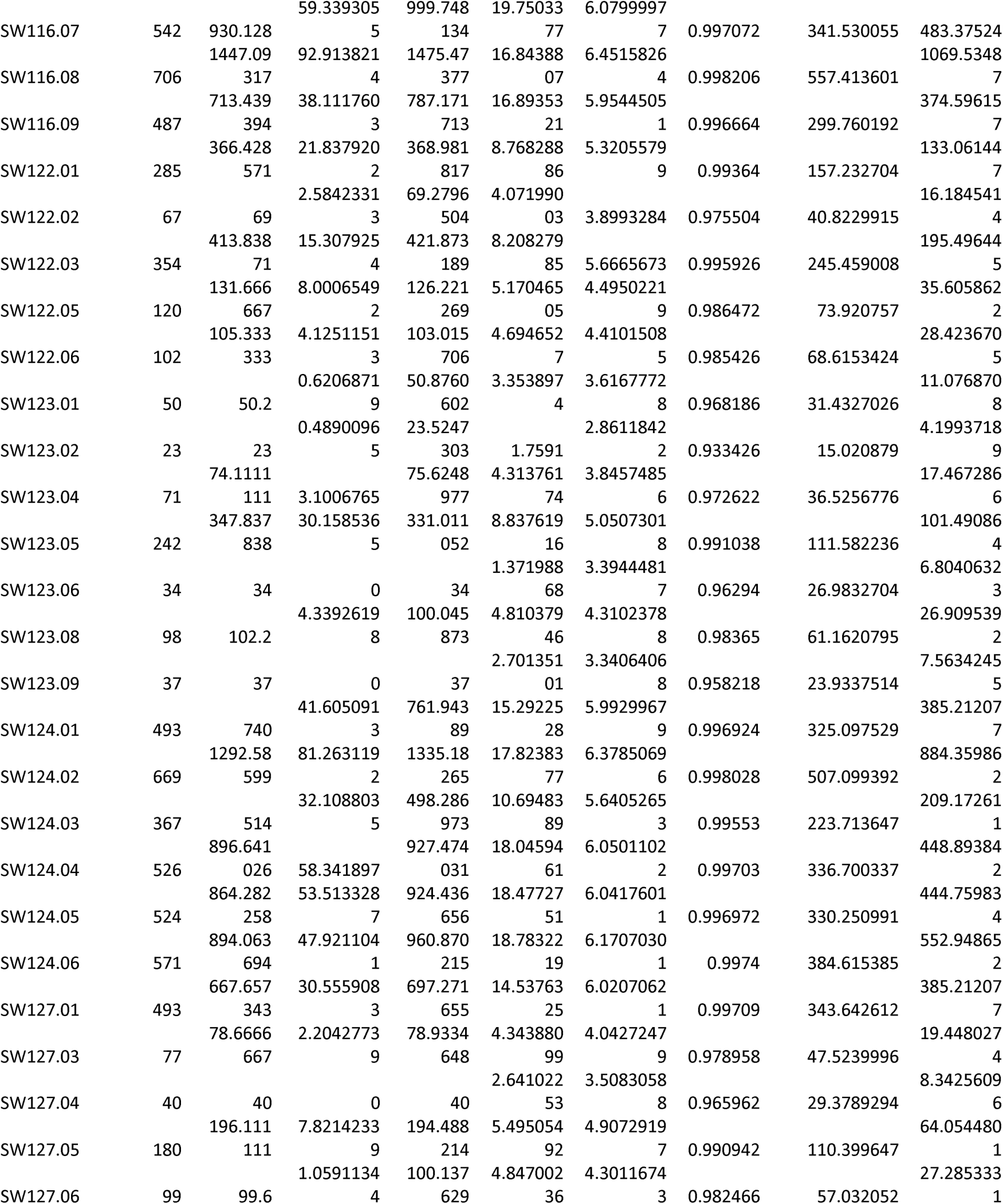
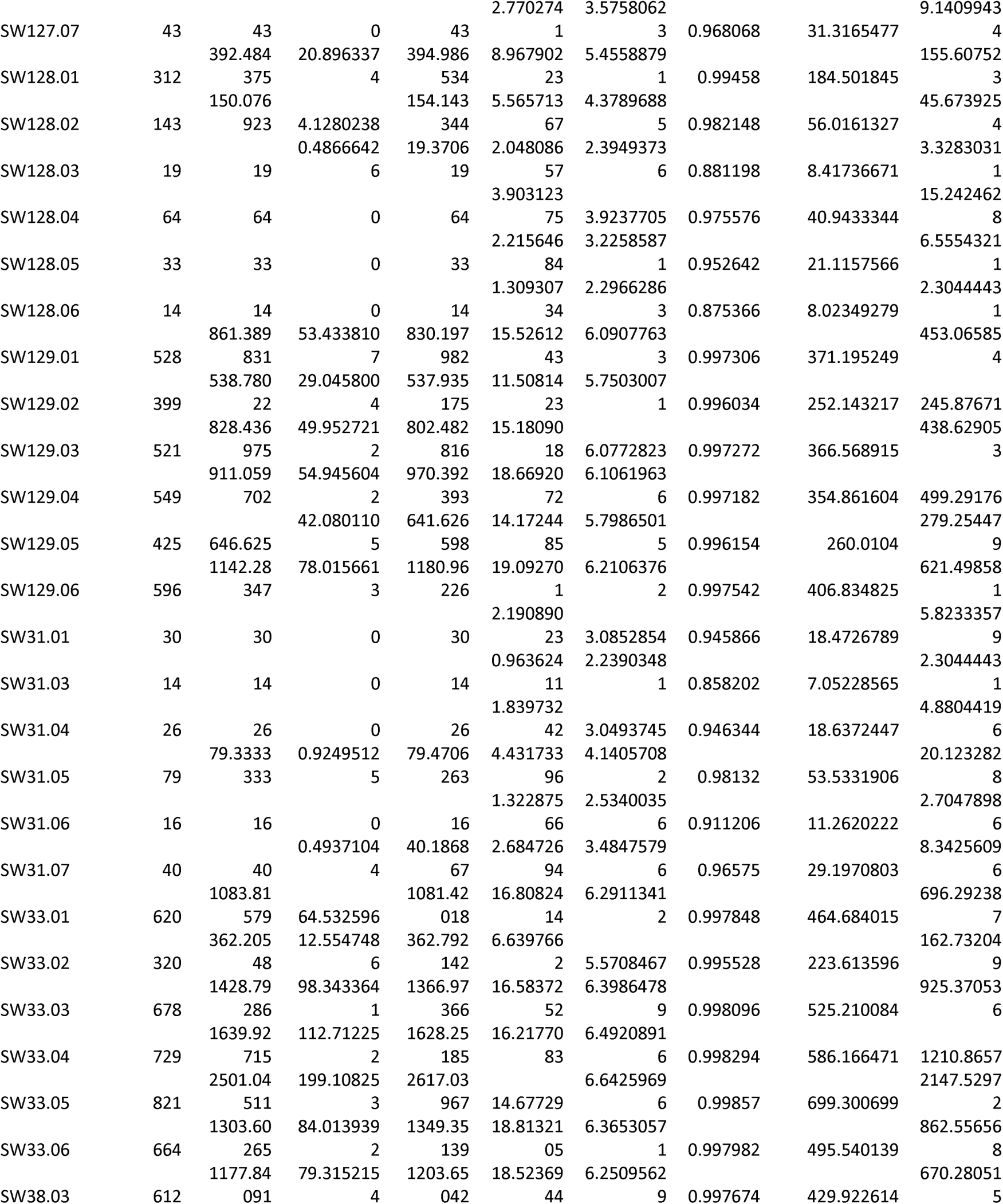
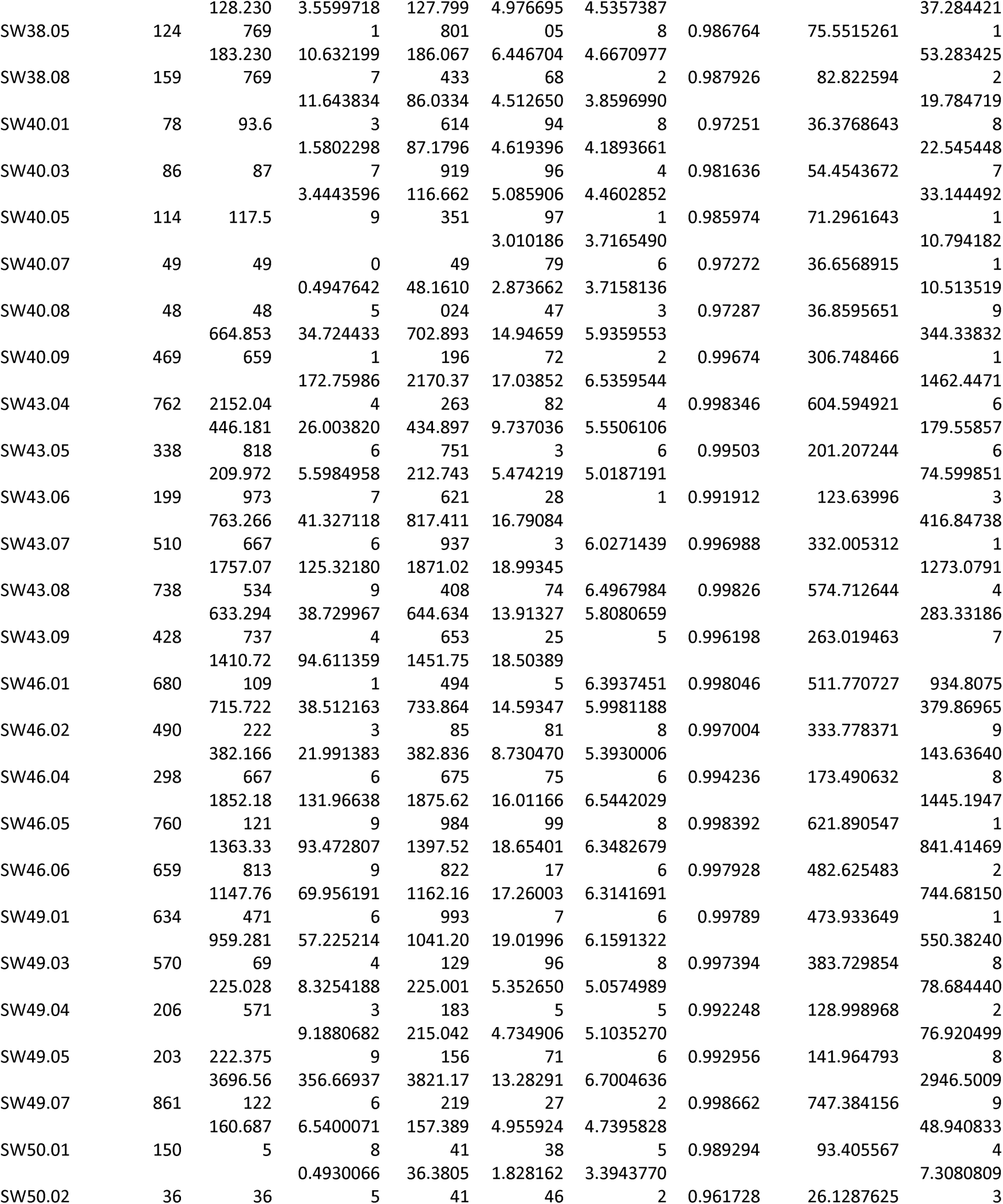
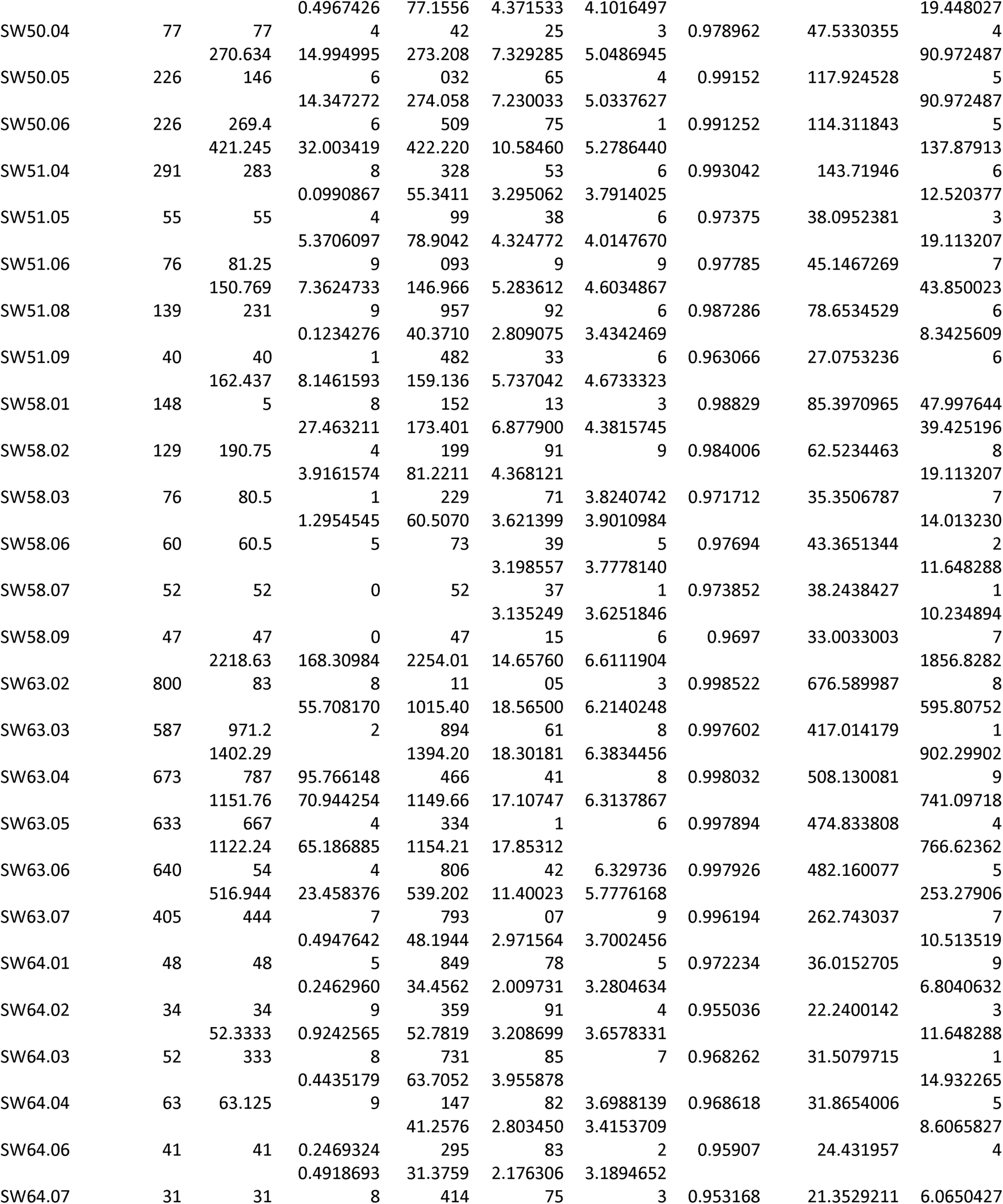
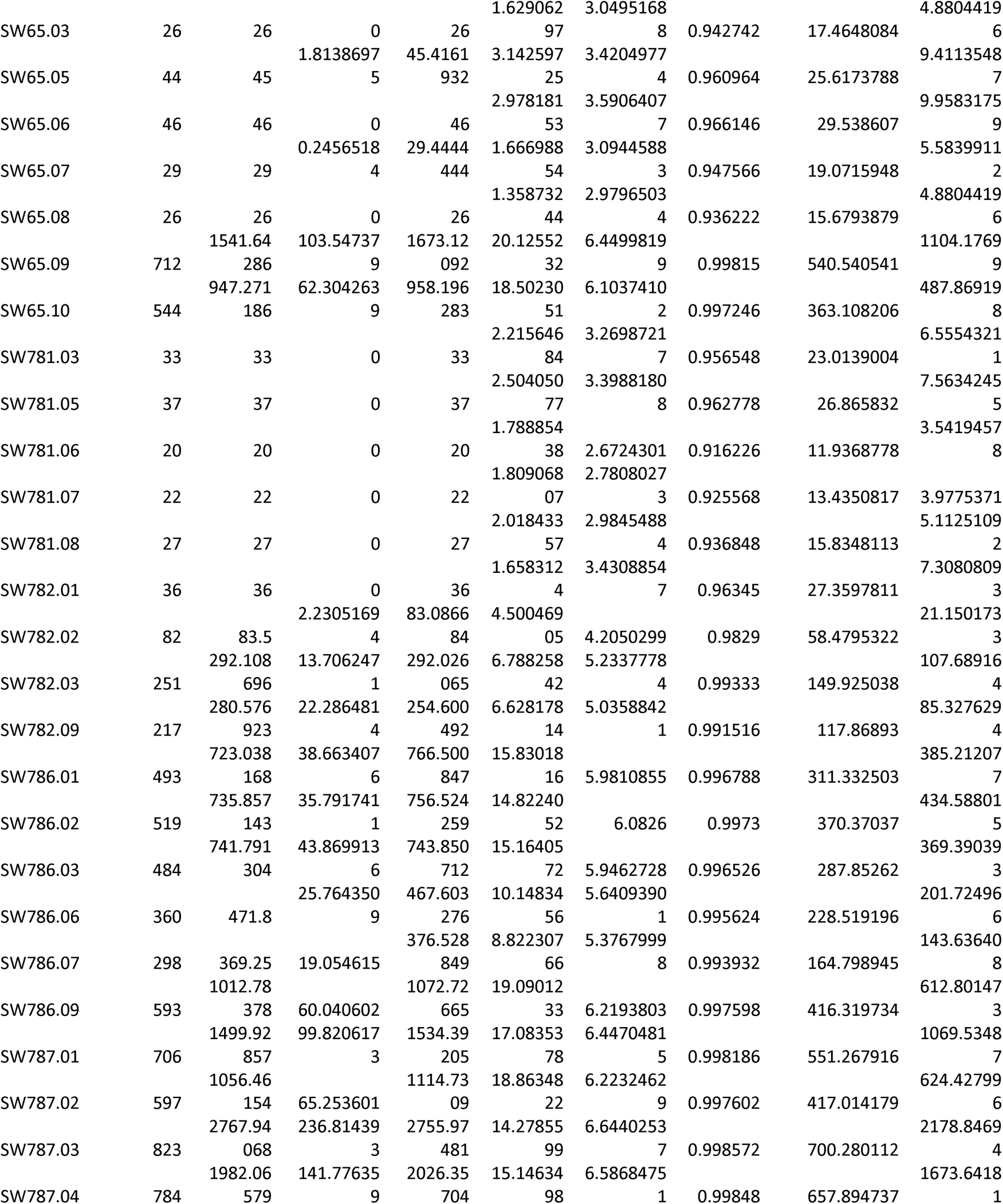
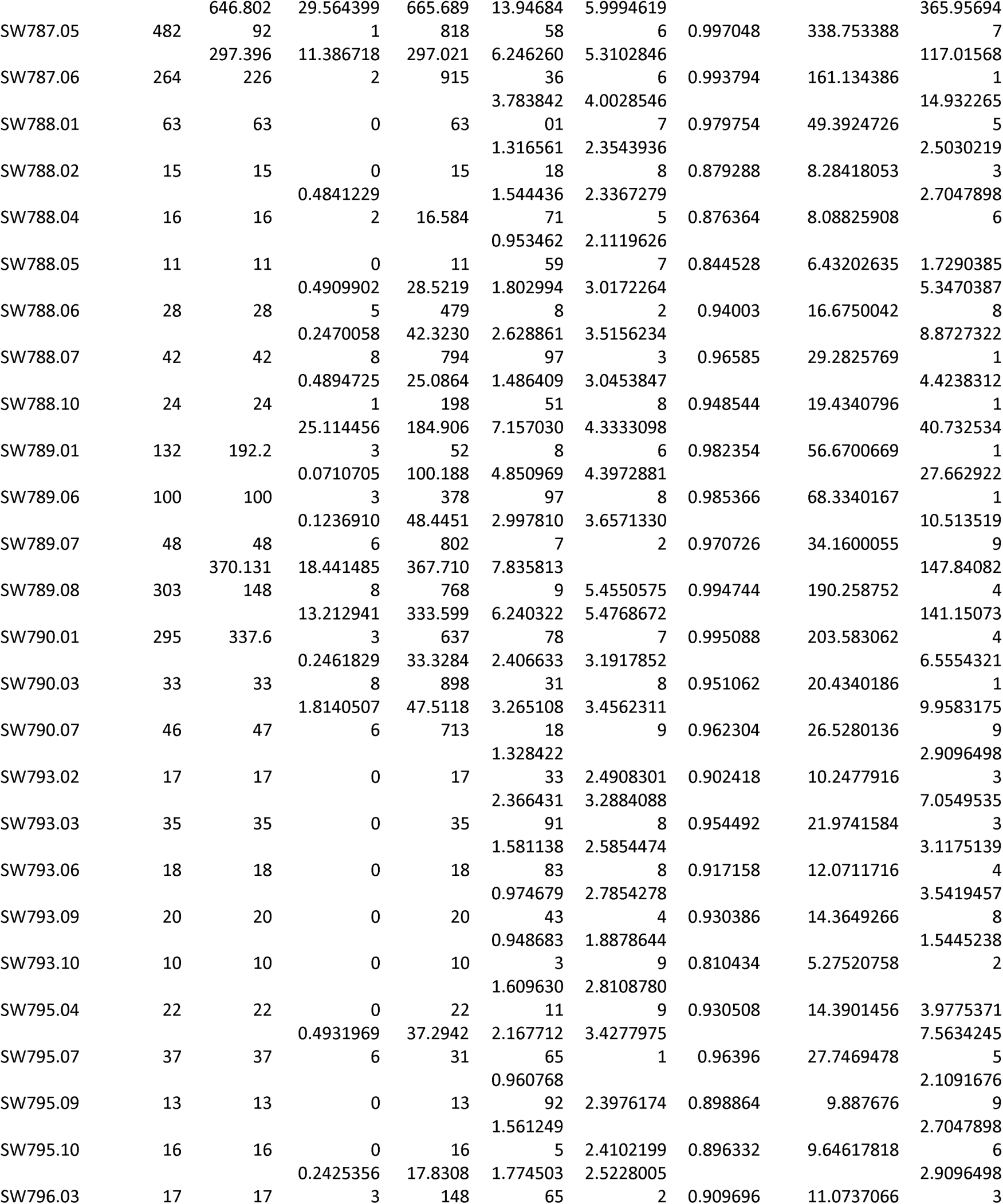
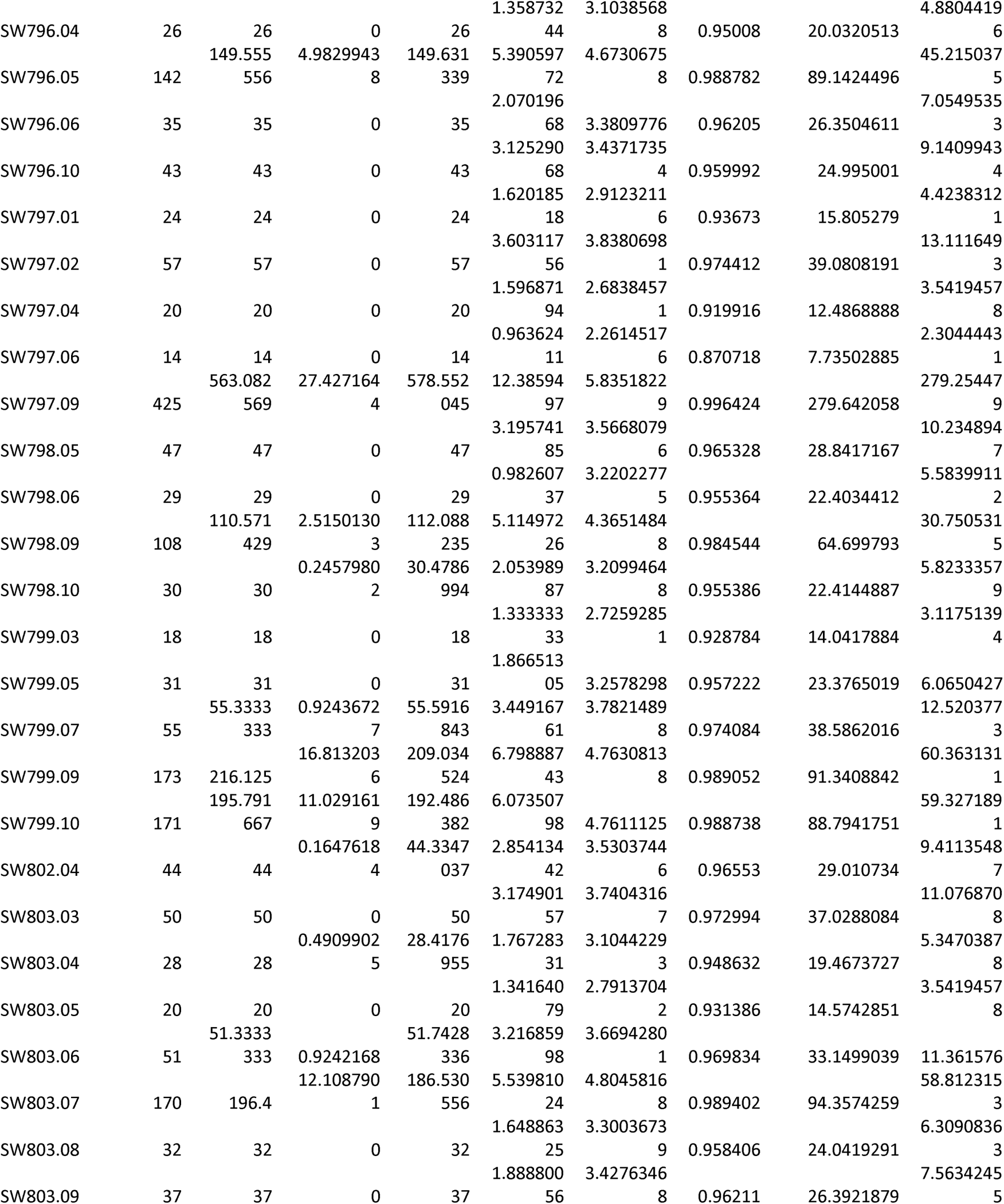
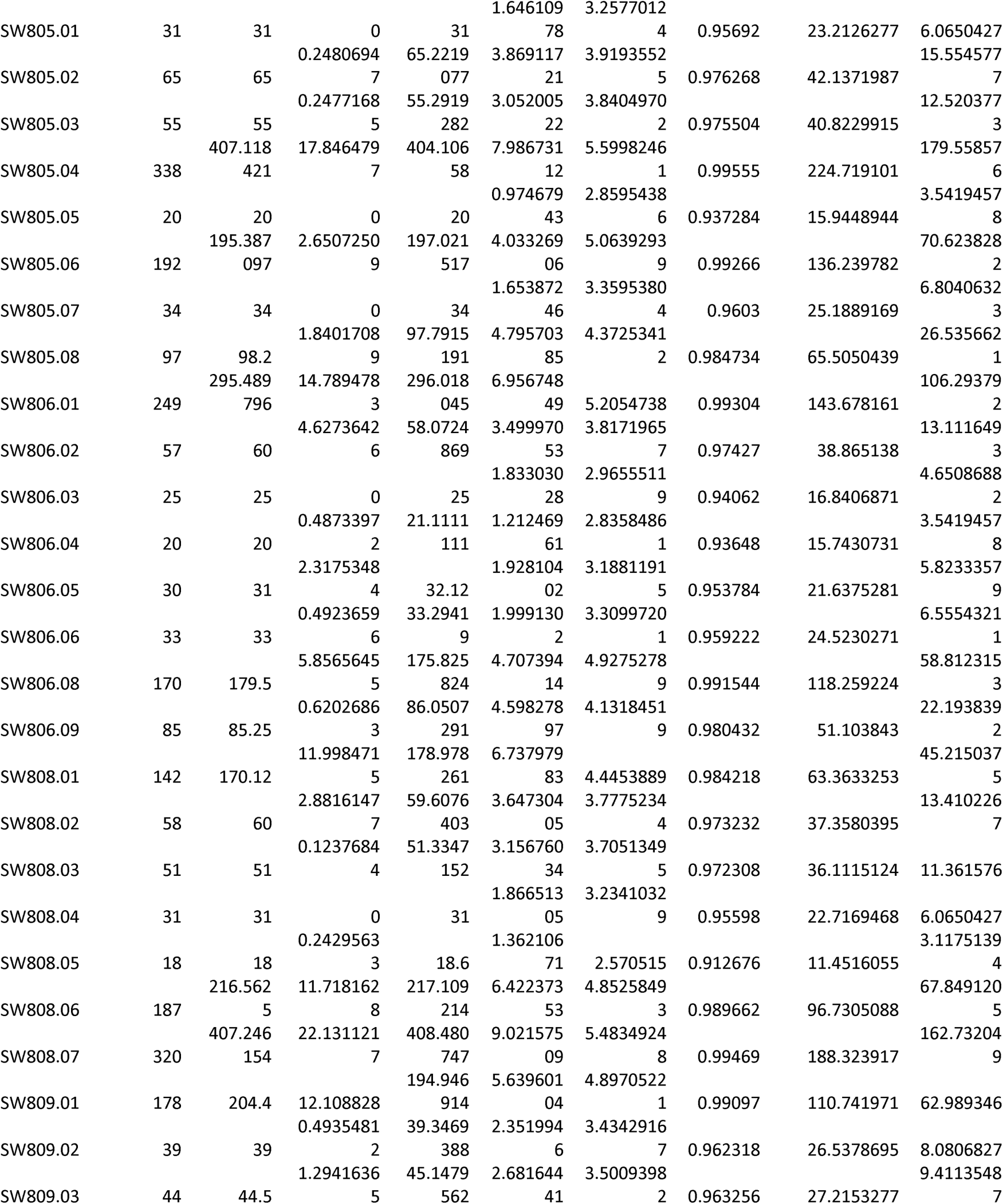
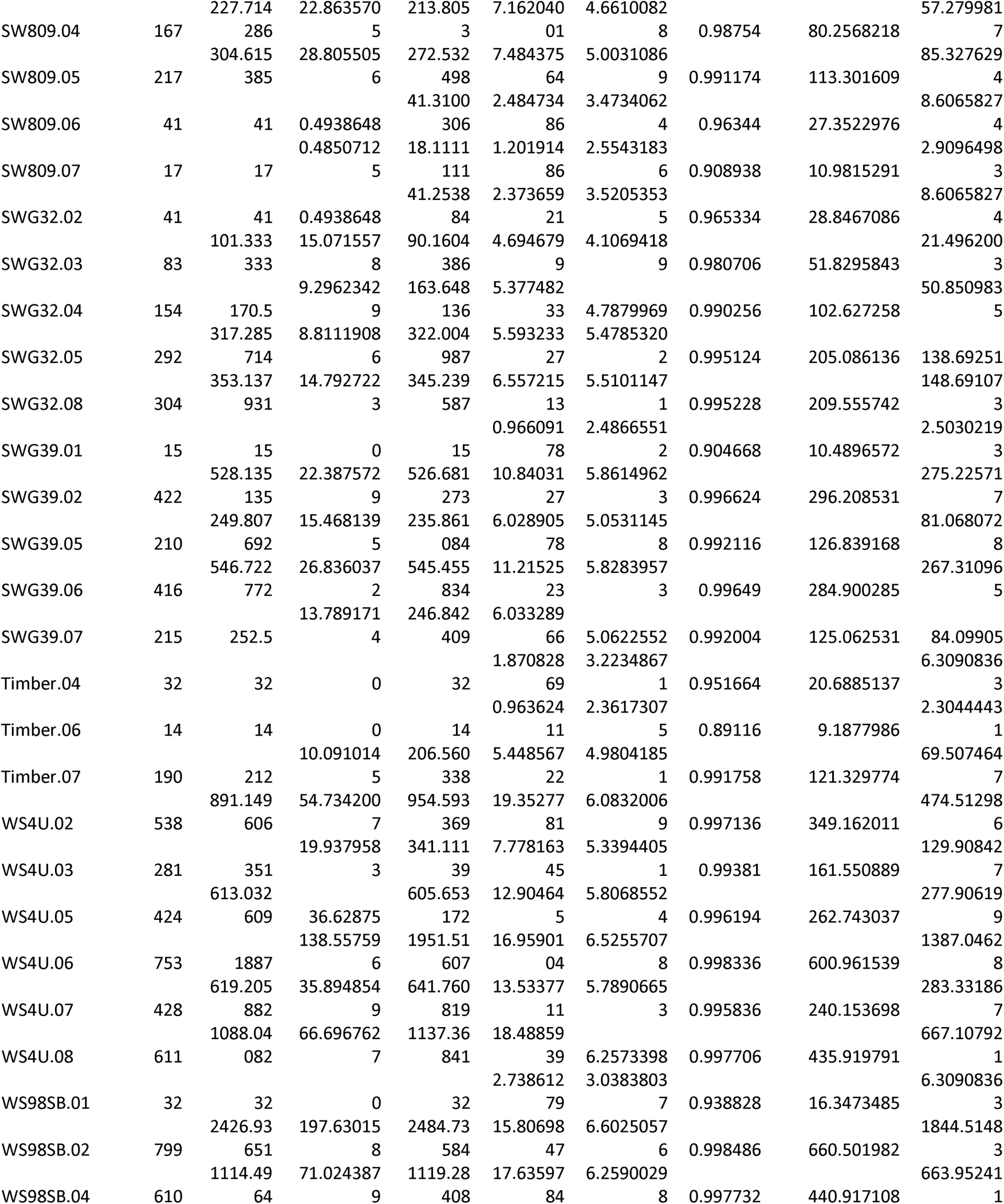
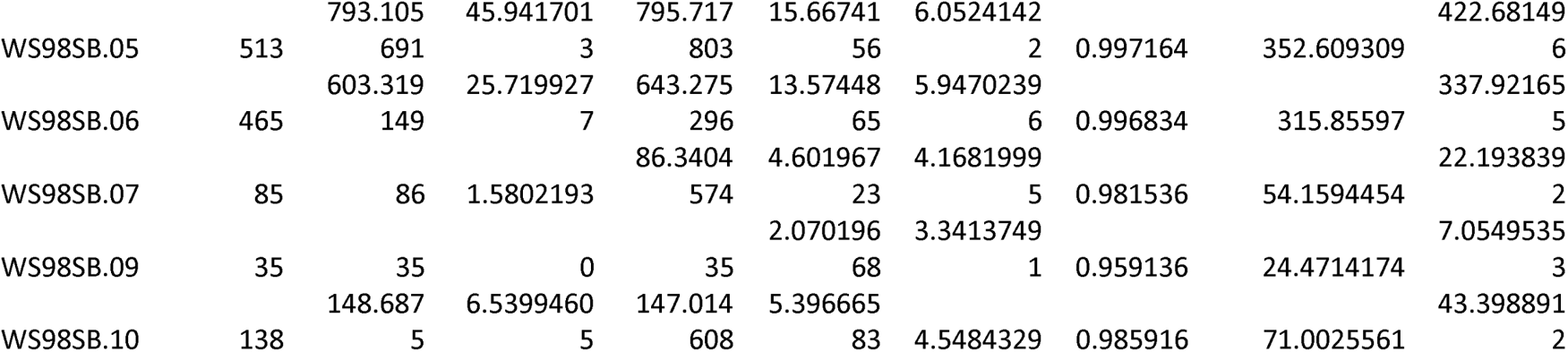
***Alpha Diversity Metrics:*** Results from the Alpha Diversity analysis for 365 switchgrass genotypes.

**Supplemental Table 2:**
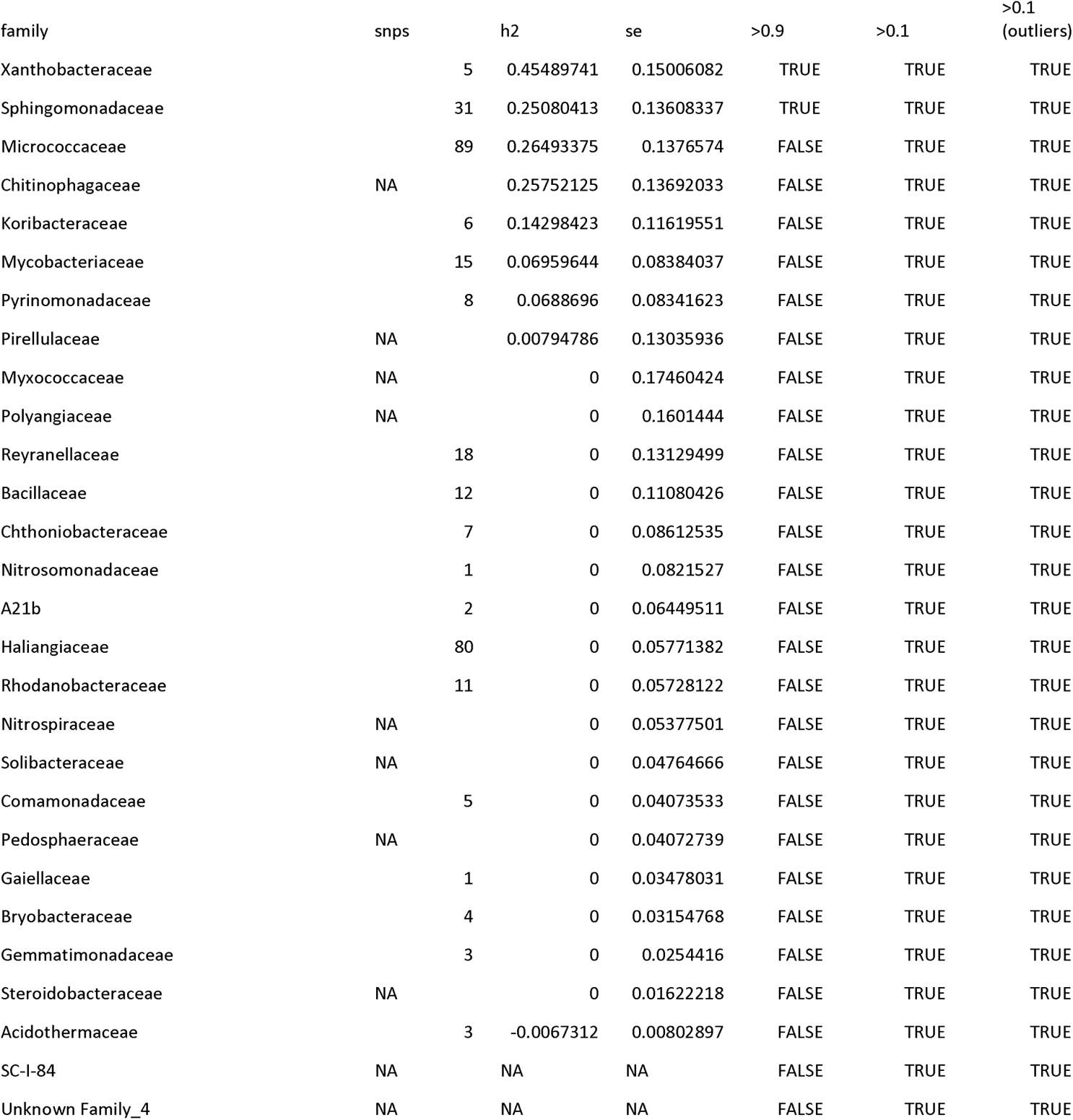
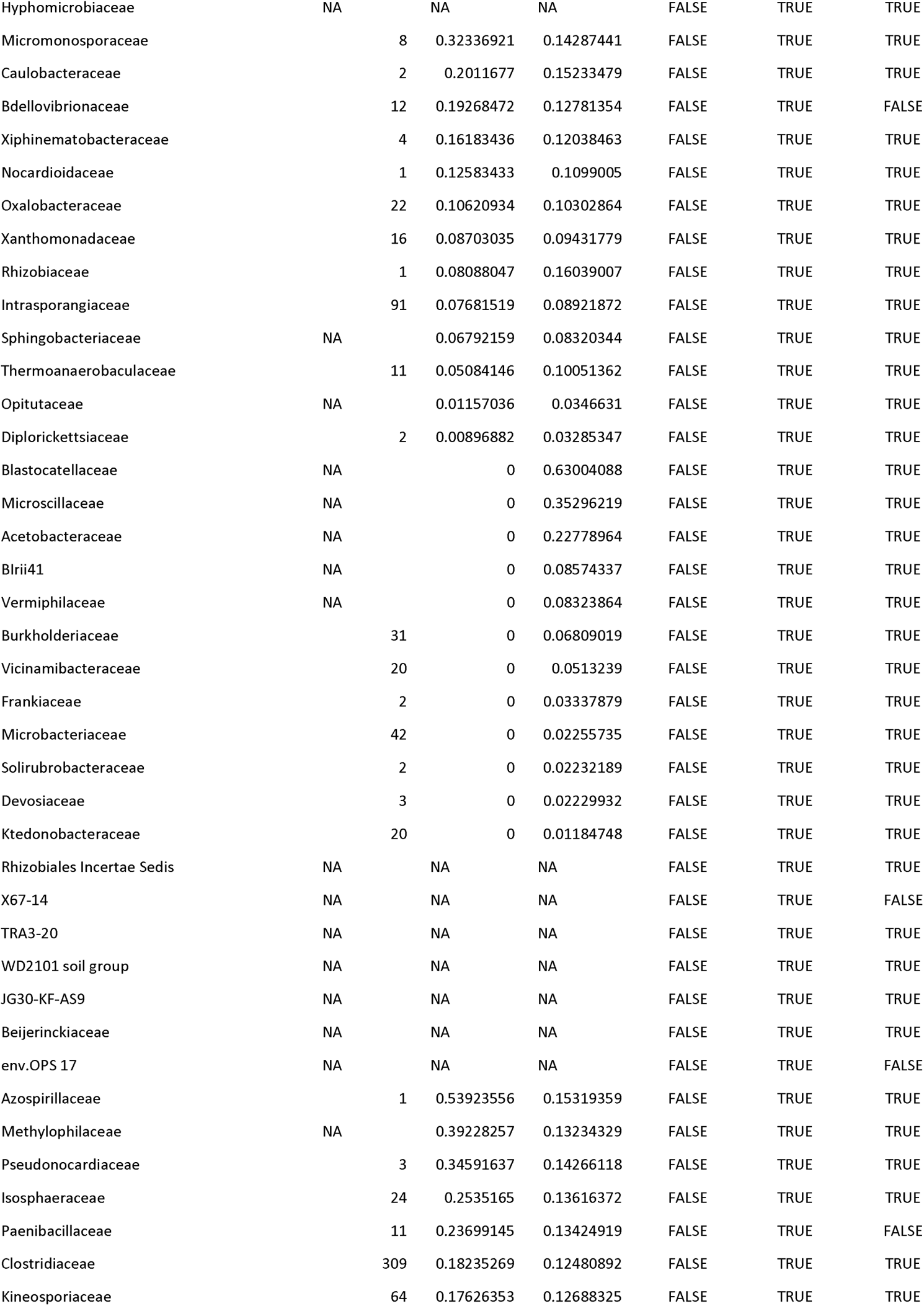
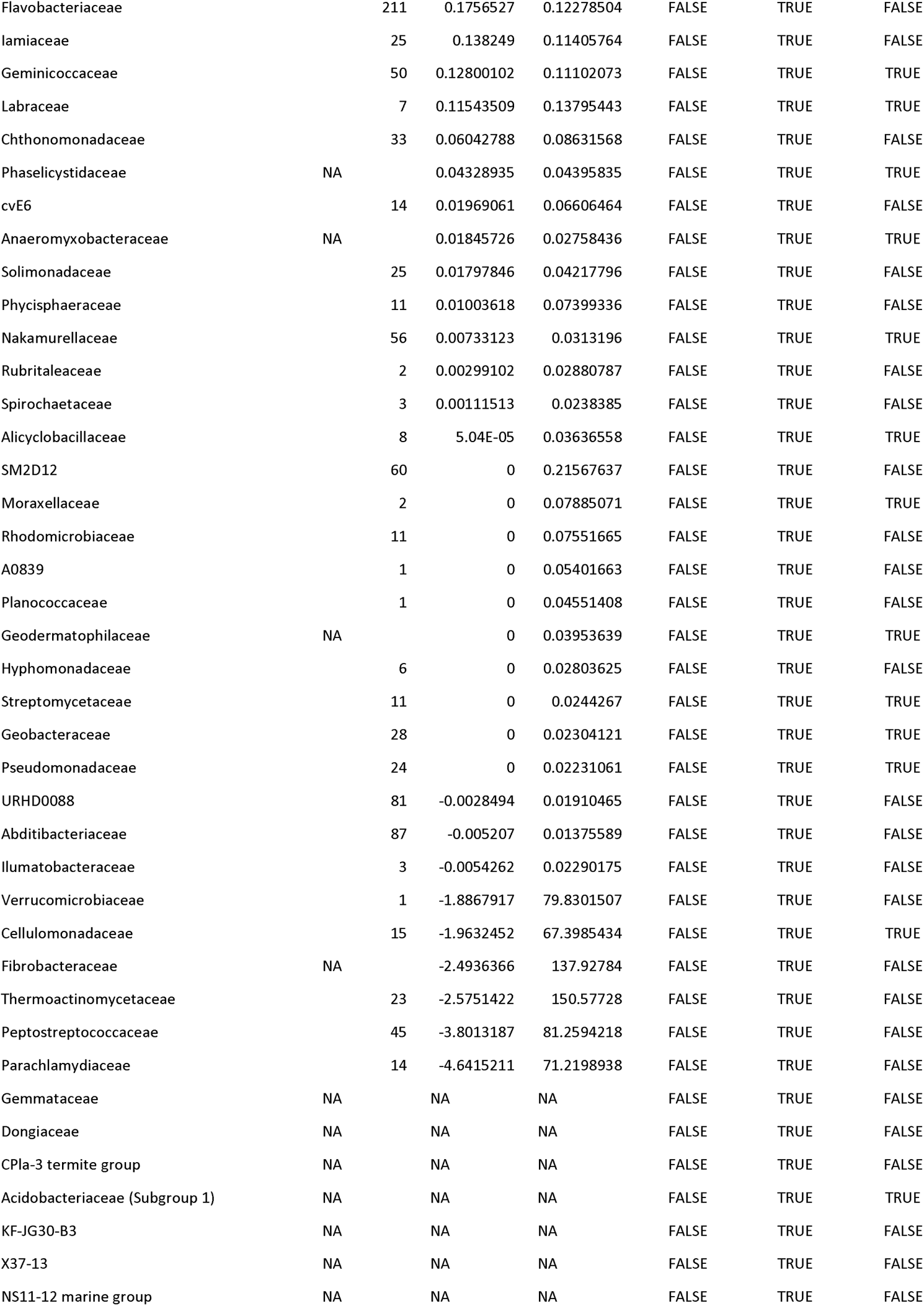
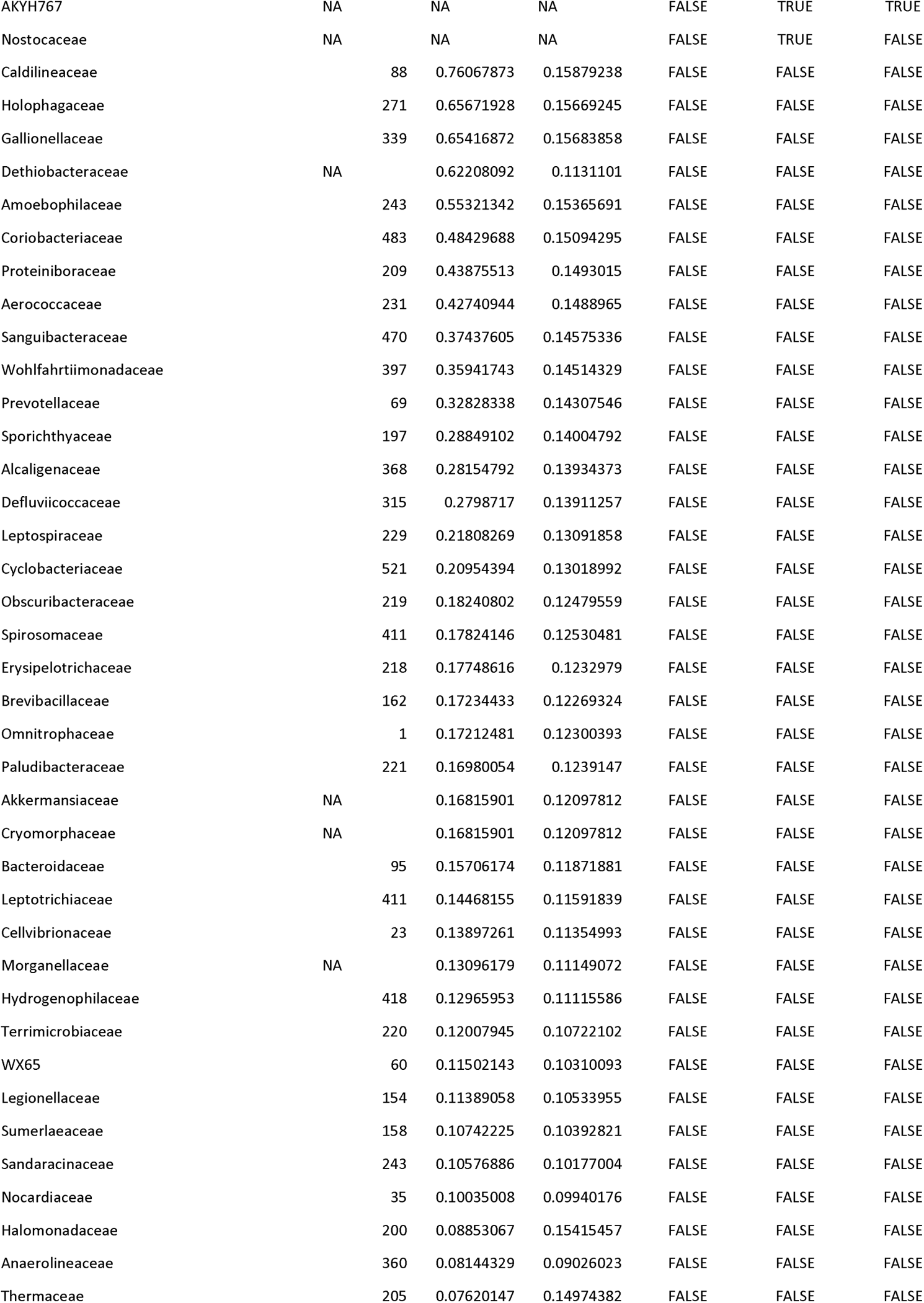
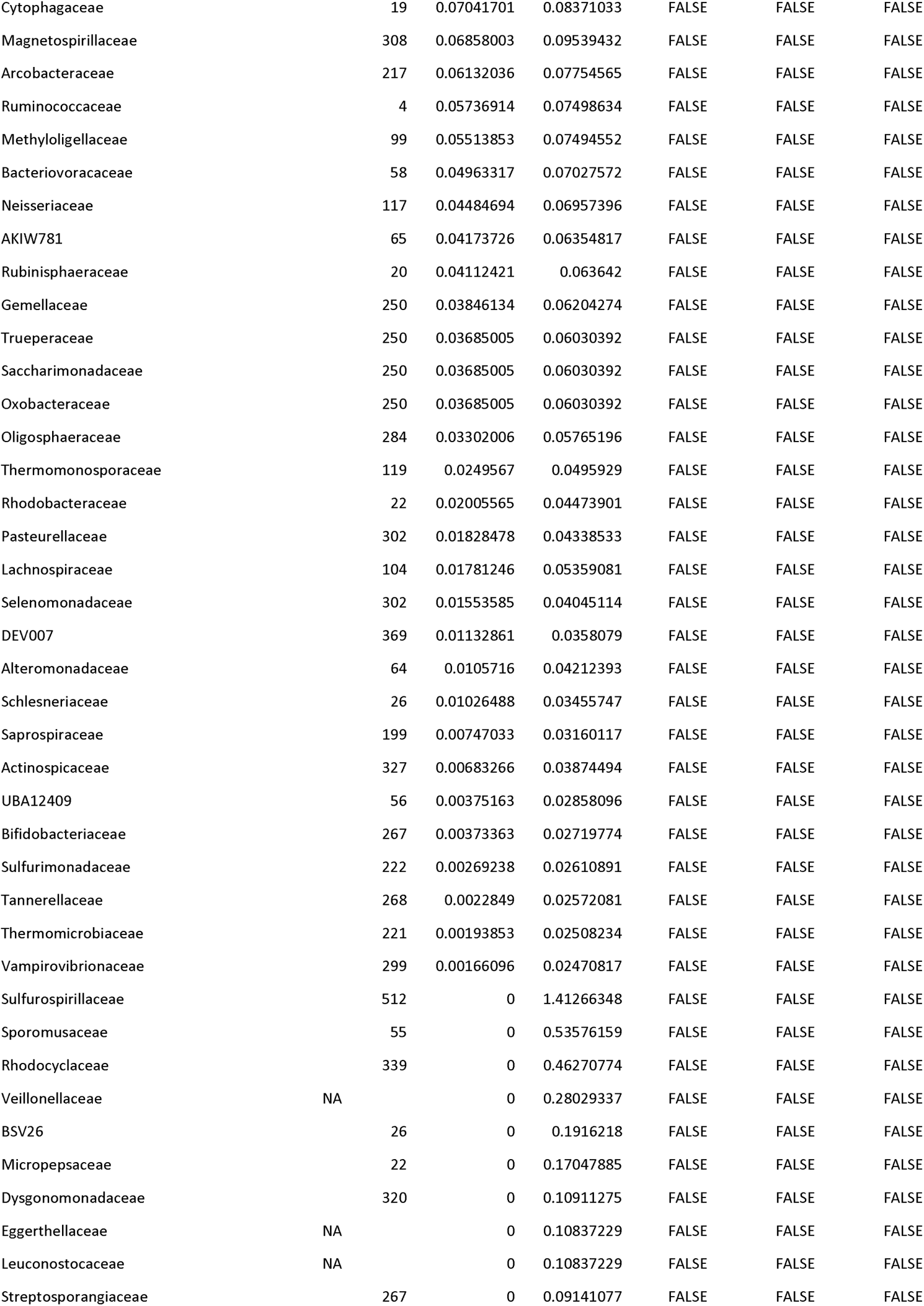
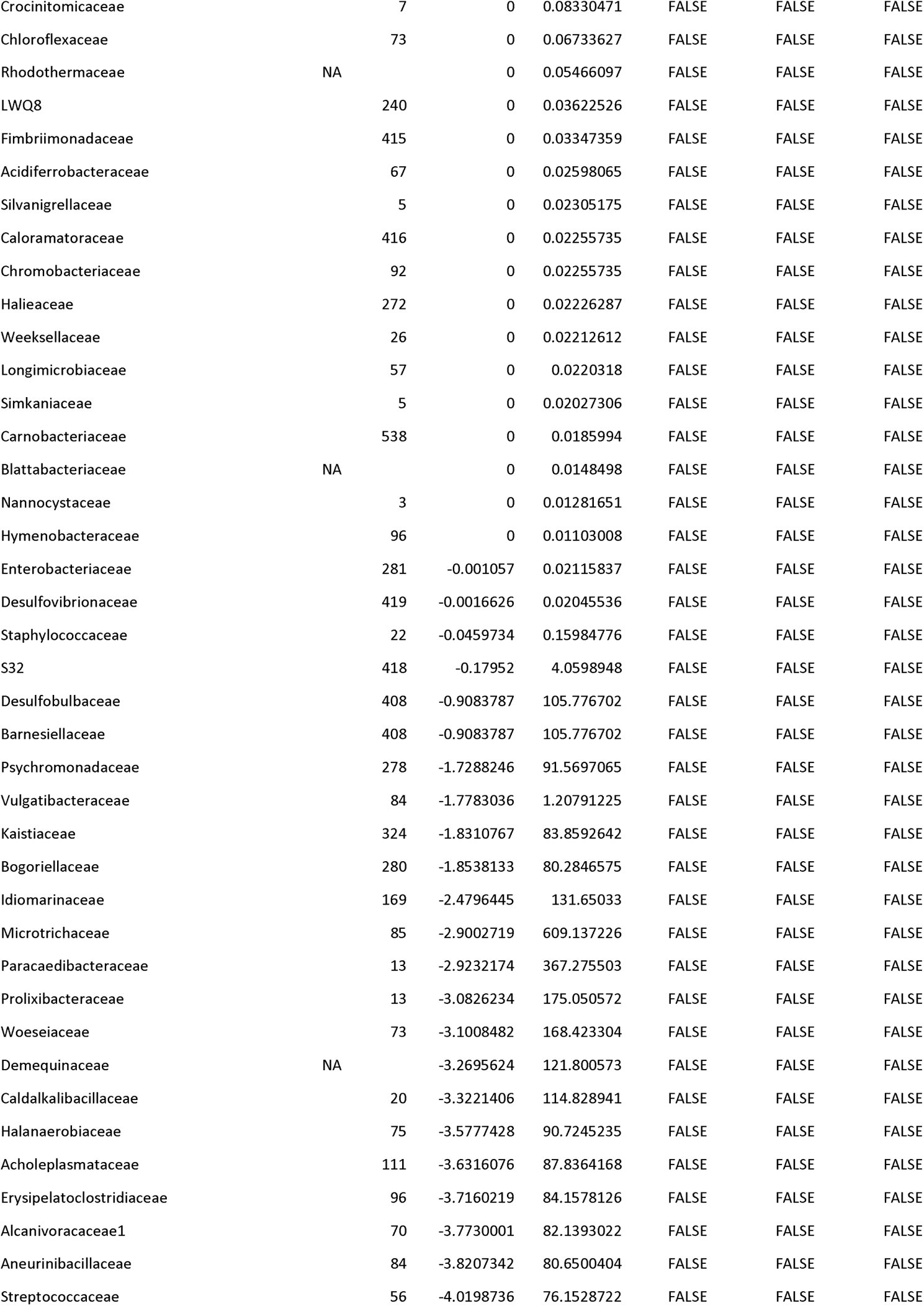
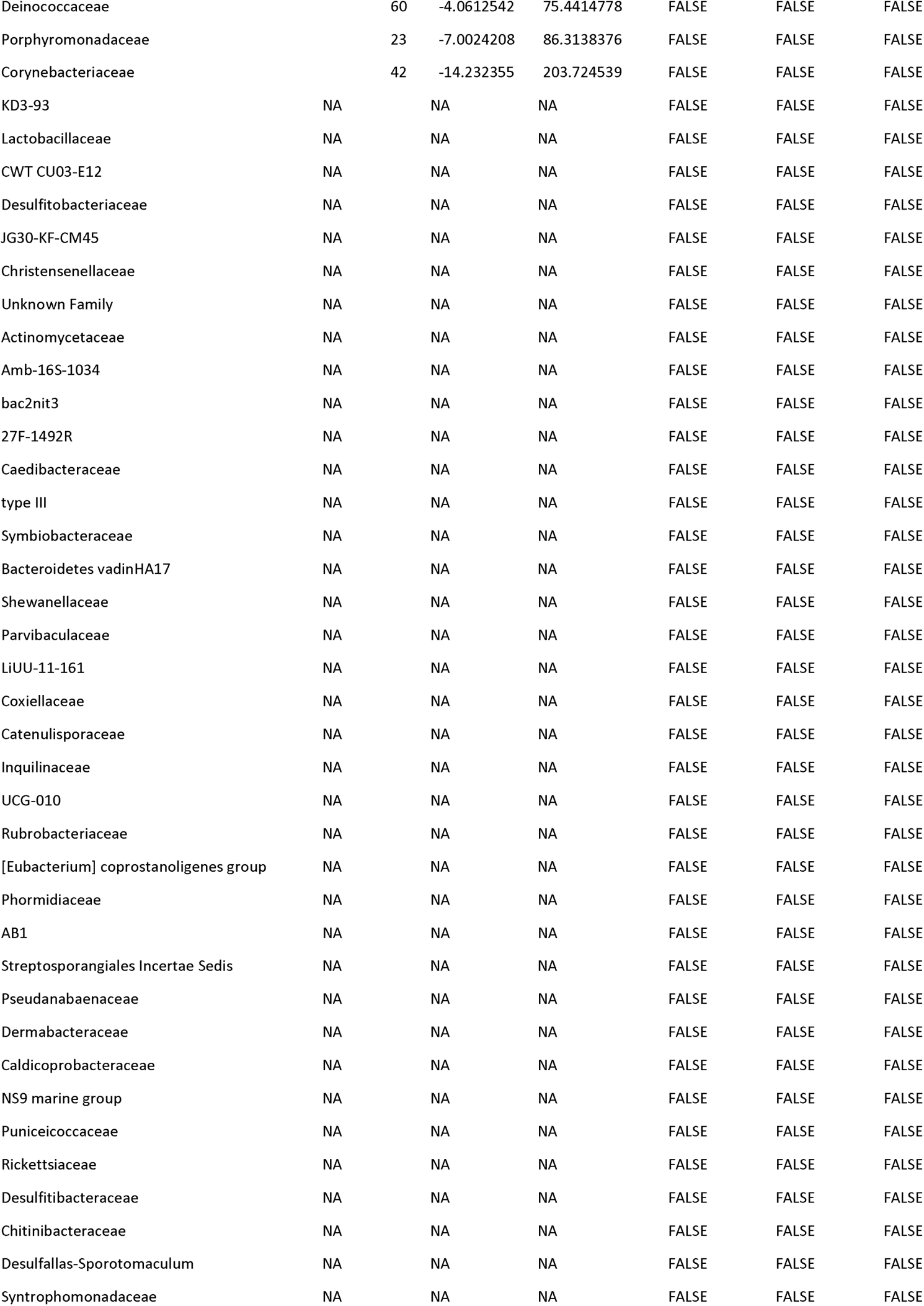
***Bacterial Family Association Data:*** Data represents the number of significant SNPs associated with each family, the heritability estimate and standard error, and an indicator for their presence above the occupancy thresholds 0.9 and 0.1 for the full ASV dataset and 0.1 for the outlier ASV dataset.

**Supplemental Table 3:**
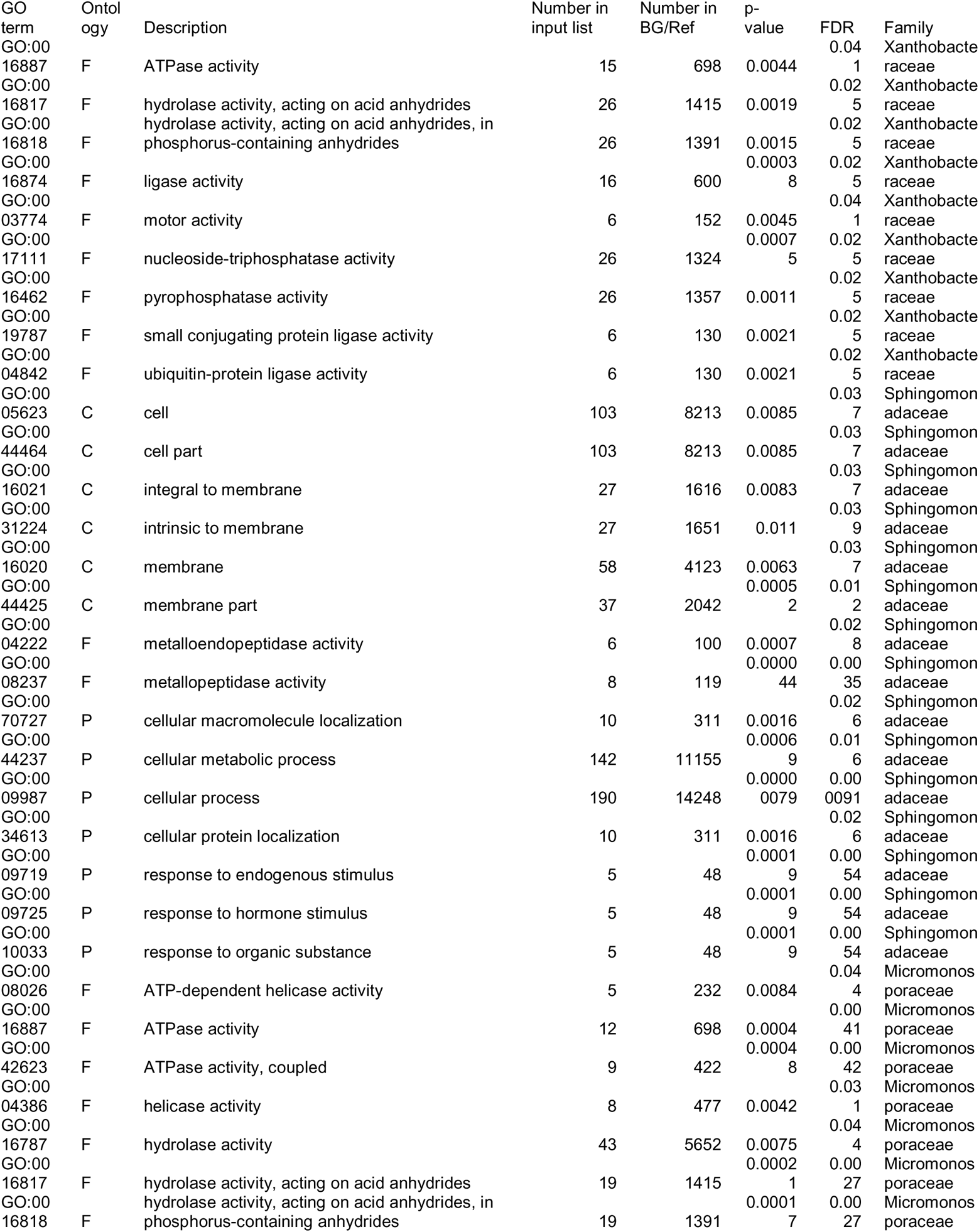
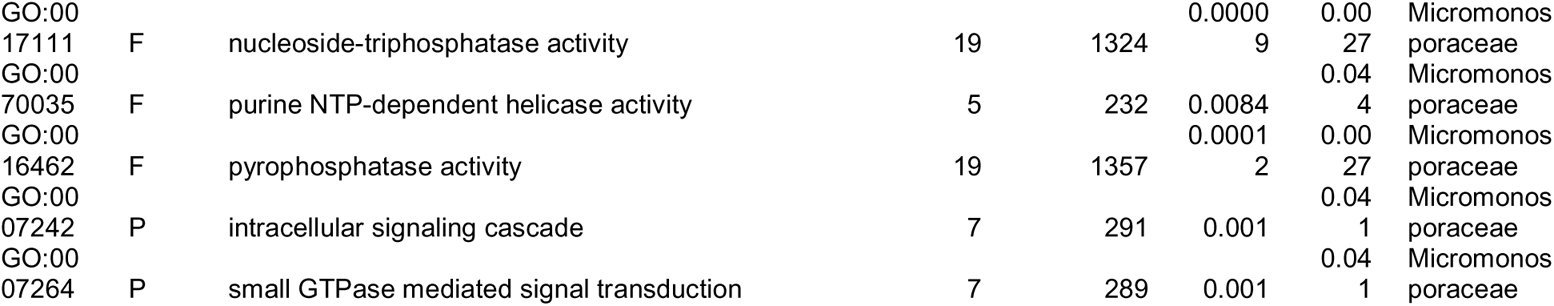
***GO Term Data:*** Data represents the Ontologies, Descriptions, Number of genes in the input list and reference genome, p-value, and FDR for GO terms associated with Xanthobacteraceae, Sphingomonadaceae, and Micromonosporaceae abundances.

